# Large-scale integration of DNA methylation and gene expression array platforms

**DOI:** 10.1101/2021.08.12.455267

**Authors:** Eva E. Lancaster, Vladimir I. Vladimirov, Brien P. Riley, Joseph W. Landry, Roxann Roberson-Nay, Timothy P. York

**Affiliations:** Department of Psychiatry, Virginia Commonwealth University, Richmond, VA 23220; Department of Psychiatry, Texas A&M University, College Station, TX, 77843; Department of Human and Molecular Genetics, Virginia Commonwealth University, Richmond, VA 23220; Department of Psychology, Virginia Commonwealth University, Richmond, VA 23220; Department of Obstetrics and Gynecology, Virginia Commonwealth University, Richmond, VA 23220

## Abstract

Epigenome-wide association studies (EWAS) aim to provide evidence that marks of DNA methylation (DNAm) have downstream consequences that can result in the development of human diseases. Although these methods have been successful in identifying DNAm patterns associated with disease states, any further characterization of etiologic mechanisms underlying disease remains elusive. This knowledge gap does not originate from a lack of DNAm-trait associations, but rather stems from study design issues that affect the interpretability of EWAS results. Despite known limitations in predicting the function of a particular CpG site, most EWAS maintain the broad assumption that altered DNAm results in a concomitant change of transcription at the most proximal gene. This study integrated DNAm and gene expression (GE) measurements in two cohorts, the Adolescent and Young Adult Twin Study (AYATS) and the Pregnancy, Race, Environment, Genes (PREG) study, to improve the understanding of epigenomic regulatory mechanisms. CpG sites associated with GE in *cis* were enriched in areas of transcription factor binding and areas of intermediate-to-low CpG density. CpG sites associated with *trans* GE were also enriched in areas of known regulatory significance, including enhancer regions. These results highlight issues with restricting DNAm-transcript annotations to small genomic intervals and question the validity of assuming a canonical *cis* DNAm-GE pathway. Based on these findings, the interpretation of EWAS results is limited in studies without multi-omic support and further research should identify genomic regions in which GE-associated DNAm is overrepresented.

## 1 Introduction

Epigenome-wide association studies (EWAS), aiming to test the theory that marks of DNA methylation (DNAm) are involved in the pathophysiology of disease, have successfully identified associations between complex traits and DNAm. ^1^ Specific DNAm patterning has been associated with environmental exposures, ^2–4^ as well as short- and long-term health outcomes. ^5–7^ Several attributes of DNAm potentially link this epigenetic mark to the development or progression of complex disease. Appropriate DNAm patterning is essential for normal development and aging, and DNAm regulatory mechanisms are implicated in a multitude of molecular processes, such as cellular differentiation, X-inactivation, and genomic imprinting. ^8–11^ As an epigenetic mark, DNAm is both dynamic and persistent; modifiable by environmental exposures yet heritable during cell division, so that any alterations to DNAm patterns may be carried through future populations of cells. ^12–15^ Importantly, altered DNAm has been linked to downstream functional changes, particularly in the regulation of gene expression (GE). These properties suggest that DNAm may be contributing to mechanisms in which previous exposures and genetic predispositions can have lasting effects on disease risk. While EWAS methods are promising, their current utility beyond biomarker discovery is questionable due to study design limitations that impact the interpretability of results, particularly those stemming from the omission of GE measurements. ^15^

The canonical mechanism describes DNAm as a repressor of proximal transcription, in which DNAm within promoter regions is able to silence GE by either blocking the binding of essential transcriptional machinery or by recruiting chromatin modifying proteins that transition the local DNA conformation to a more heterochromatic state. ^10,16^ Despite accumulating evidence that suggests this model is overly simplistic, many researchers rely on this paradigm to interpret an association between DNAm and a disease of interest. In a typical EWAS, any significantly associated CpG sites (also known as differentially methylated positions or DMPs) are each mapped to their most proximal gene and the biological function of those genes is reported in the context of the tested phenotype. Given that this interpretation emphasizes the functional relevance of specific genes to disease biology, an argument can be made that current EWAS are primarily interested in examining a theory of DNAm-driven transcriptional regulation. ^15^ By inferring transcriptional activity from DNAm-trait associations, this approach relies on assumptions without directly testing for functional evidence. Given that accurately inferring the functional consequences of modified DNAm at any particular site is still very limited, this practice may lead to inaccurate conclusions about disease biology. ^17^

Accurately predicting the functional impacts of altered DNAm remains challenging, in part, due to the limited characterization of genome-wide DNAm-GE relationships. ^18^ DNAm often does not block transcription independently but rather works in concert with other regulatory elements to coordinate GE. These regulatory mechanisms involve a complex crosstalk between DNAm, higher-order chromatin modifiers, and other epigenetic marks, further contributing to difficulties in determining the functional impact from DNAm measurements alone. ^19–21^ Moreover, linking genes to their putative regulatory regions is not always straightforward. ^22^ DMPs are often located outside of proximal regulatory elements, within intergenic or intronic regions with no known regulatory function. Since a frequently utilized approach for interpreting these results involves linking all DMPs to their nearest gene, any features of the genomic landscape beyond distance are disregarded. Even if CpG-GE pairs are identified, predicting the regulatory consequence of altered DNAm remains difficult as exceptions to the canonical theory have accumulated. For example, increased DNAm, particularly within the gene body, is frequently positively correlated with local transcription. ^23–28^ Although mechanisms linking hypermethylation to increased GE are still unclear, a recent study identified more transcription factors which preferred binding methylated sequences than those inhibited by DNAm. ^21^ These functional complexities suggest that assumptions regarding transcriptional activity should not be inferred by DNAm patterns alone. Instead, if the fundamental theory being explored is a mechanism of transcriptional regulation modulated by DNAm, measurements of GE should be included in the analysis. ^15^

Multi-omic studies integrating global DNAm and GE measurements can provide evidence for DNAm-driven transcriptional regulatory mechanisms. Measuring GE alongside DNAm would allow for direct testing of the proposed mechanism while avoiding assumptions regarding the regulatory function of DNAm that typically cloud the interpretation of EWAS results. In order to model the proposed molecular mechanism, analyses testing DNAm-driven transcriptional regulation should provide evidence for the mediating effect of GE on the phenotype of interest. Instead, studies often discover differentially methylated and differentially expressed genes separately by performing both an EWAS and transcriptome-wide association study, and any overlaps between DNAm-trait and GE-trait associations are reported. ^29–36^ While the addition of GE-trait associations provides more support for functional changes within the cell, it is unclear what hypothesized biological mechanism this analysis is testing. Moreover, since the relationship between DNAm and GE is never tested, this method still relies on assumptions that DMPs strictly influence expression of the nearest transcript.

An extended EWAS approach integrating both DNAm and GE measurements holds promise in uncovering biological processes important to the development or progression of disease, however, a mechanistic interpretation requires prior knowledge regarding specific DNAm-GE relationships across the genome, which has yet to be resolved. The objective of this study was to catalogue the relationships between DNAm and both proximal and distal GE (i.e., *cis* and *trans* relationships, respectively) in peripheral blood, a tissue commonly assayed in EWAS. To identify attributes that replicate across disparate samples, analyses were conducted in two previously described cohorts, the Adolescent and Young Adult Twin Study (AYATS), ^7,37^ and the Pregnancy, Race, Environment, Genes Study (PREG). ^38^ An in-depth characterization of GE-associated CpG sites could improve predictions of the downstream functional impact of altered DNAm and inform best practices for interpreting DNAm-trait associations generated by EWAS.

## 2 Methods

### 2.1 Study cohorts

#### Adolescent and Young Adult Twin Study (AYATS)

The AYATS study was designed to examine genetic and environmental contributions to internalizing pathways (e.g., depression and anxiety) during development. A sample of monozygotic twins were chosen for their adherence to the study’s inclusion criteria (e.g., 15-20 years of age, no current use of psychotropic medications). ^7,37^ Peripheral blood collected from 141 participants at a single time point was assayed for both DNAm and GE. An overview of study characteristics and further demographic information can be accessed in the supplement (Supplementary Table S1).

#### Pregnancy, Race, Environment, Genes (PREG) study

The PREG Study is a prospective longitudinal study with the purpose of identifying how environmental determinants of health and DNAm remodeling relate to racial health disparities in perinatal health outcomes. ^38^ Of the 240 women who enrolled in the study, 177 met all birth and pregnancy inclusion criteria (e.g., mother and father self-identify as either both Caucasian or both African American) and no exclusion criteria (e.g., preeclampsia, fetal congenital anomaly, placental anomaly, fewer than 3 study time points completed). Peripheral blood samples were collected up to four times throughout pregnancy. Sample collection was scheduled during gestational weeks 0-15, 10-25, 20-40, and 37-42. DNAm was assessed at all time points whereas GE was measured once at the final collection during weeks 37–42. Only those DNAm measurements from specimen simultaneously collected with GE were analyzed in this study. A total of 151 women had concomitant DNAm and GE measured. An overview of study characteristics and further demographic information can be found in the supplement (Supplementary Table S2).

### 2.2 DNAm measurement and data processing

In both samples, DNAm and GE was measured from peripheral blood. The Infinium 450k HumanMethylation BeadChip assayed genome-wide DNAm and the Affymetrix HG-U133A 2.0 array measured GE. A description of platform characteristics as well as the methods used for measurement and preprocessing can be found in the supplement.

### 2.3 Association analysis

The relationship between all pairwise combinations of measured DNAm and GE (Table 1) was tested by linear regression in the R statistical environment (version 3.5). ^39^ Log-transformed expression values (dependent variable) were regressed on DNAm M-values (independent variable), while covariates controlled for differences in cell type heterogeneity. Cell type proportions were derived from the Houseman algorithm, which estimates proportions for granulocytes, monocytes, CD8-positive T cells, CD4-positive T cells, B lymphocytes, and natural killer cells based on cell type-specific DNAm profiles. ^40^ Granulocytes were selected to account for overall differences in cell type proportions based on high correlations with other cell type estimates (absolute correlations ranged from 0.47 to 0.71), and included as a covariate in all models. Natural killer proportions were included in AYATS models exclusively to adjust for the atypical variation in this cellular fraction characteristic of depressed patients. ^7,41^

**Table 1:**
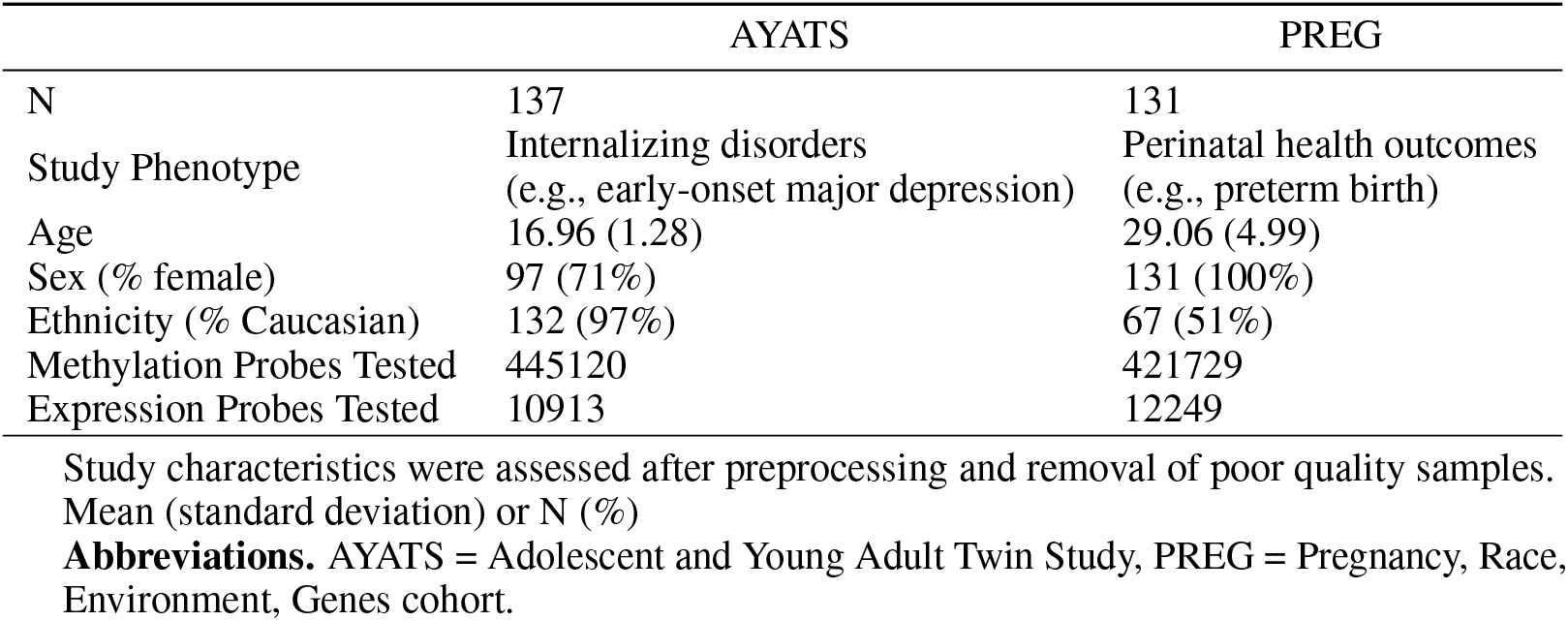
Study Characteristics

|  | AYATS | PREG |
| --- | --- | --- |
| N | 137 | 131 |
| Study Phenotype | Internalizing disorders<br>(e.g., early-onset major depression) | Perinatal health outcomes<br>(e.g., preterm birth) |
| Age | 16.96 (1.28) | 29.06 (4.99) |
| Sex (% female) | 97 (71%) | 131 (100%) |
| Ethnicity (% Caucasian) | 132 (97%) | 67 (51%) |
| Methylation Probes Tested | 445120 | 421729 |
| Expression Probes Tested | 10913 | 12249 |

Study characteristics were assessed after preprocessing and removal of poor quality samples. Mean (standard deviation) or N (%)
**Abbreviations.** AYATS = Adolescent and Young Adult Twin Study, PREG = Pregnancy, Race, Environment, Genes cohort.

Additional covariates were selected to adjust for potential confounding influences specific to the characteristics of each cohort, while also maintaining a similar analytical approach across the two studies. Since PREG is a racially diverse sample (Table 1), ancestrally informative principal components were estimated from the DNAm data using the method described in Barfield et al. The third principal component was highly correlated with self-reported race and included as a covariate in the PREG cohort models. ^42^ A linear mixed-model framework was used to account for twin structure in the AYATS cohort. ^43^ The limma Bioconductor package was used to estimate within-family correlations from 1,000 randomly sampled CpGs in order to appropriately adjust model standard errors and account for the non-independence of twin pair DNAm observations. ^43^

Both DNAm and GE measurements were adjusted for technical artifacts prior to analysis (see supplement), so that variables related to slide or row effects were not included as covariates in subsequent analyses.

Although measurements were generated using the same technology in both cohorts, differing numbers of probes remained after quality control procedures (Table 1). A within-study Bonferroni correction was used to adjust for multiple testing at an alpha threshold of 0.05. While estimates of genomic inflation are typically used to identify spurious associations driven by artefacts in genome-wide association studies (GWAS), it has been recently suggested that inflated test statistics should be similarly reviewed in epigenetic studies. ^44^ To mitigate the presence of false positives, genomic inflation was assessed using the method described in Kennedy et al. ^24^ Briefly, genomic inflation factors were calculated for each transcript, across all CpG associations, as the median (T-statistic)^2^/0.4549. Appropriate thresholds for test statistic inflation are not as well established in the epigenetics field. To facilitate cross-study comparisons, any transcript with an inflation factor > 2 was flagged for removal. ^24^

Every pairwise relationship between measured DNAm and GE was modeled and classified as either *cis* or *trans*, since molecular mechanisms linking proximal DNAm may differ from more long-range interactions. DNAm-GE pairs were in *cis* if the CpG site was located within a gene or 2,500 base pairs upstream. This extension is expected to capture important transcript-specific regulatory regions, given that many promoters are located up to 1 kilobase upstream of the transcriptional start site (TSS). ^45^ CpG-transcript pairs located outside this range were categorized as *trans* relationships, with the rationale that more distal regulatory features (e.g., enhancers) may be responsible for the relationship between CpG methylation and transcript expression.

### 2.4 Characterization of results

CpG sites were mapped to biologically relevant annotations to test for feature enrichment among sites significantly associated with GE. Annotation selection was based on evidence that local CpG density, gene feature location, and proximity to regulatory elements are important characteristics that may impact the functional consequences of DNAm. ^16^ Transcription factor activity is regulated by DNAm, and a history of transcription factor binding also appears to influence the susceptibility of CpG methylation in specific locations. ^21,46–48^ Moreover, processes involving transcription factors were previously enriched among CpGs associated with GE. ^24,49^ Similarly, non-coding RNAs are regulated by methylation patterning, while also contributing to the regulatory activity of DNAm. ^50,51^ Chromatin states are accurately able to distinguish variable transcriptional activity by describing the specific patterns of histone modifications that impact the regulation of GE. ^52,53^ Histone modifications are intricately linked to both DNAm and GE, potentially serving to mediate the influence of methylation on transcription. ^16,52^

Selected features described local CpG densities (UCSC CpG island classifiers and HIL annotations), ^54,55^ genomic regions, chromatin states (ENCODE 15-state ChromHMM), ^53^ transcription factor binding (ENCODE TF ChIP-seq), ^56^ non-coding RNAs (GENCODE version 37), ^57^ and other annotations related to regulatory activity (i.e., FANTOM5-defined enhancer and ENCODE-defined insulator regions). ^58^ All cell type-specific annotations (e.g., chromatin states, enhancer regions, etc.) were defined in the lymphoblastoid cell line GM12878.

Enrichment analyses were performed separately for *cis* and *trans* groups. The proportion of significant findings annotated to each category was compared to the proportion of total number of tested CpG sites using Fisher’s exact test. A Bonferroni correction for 20 enrichment tests was used to adjust for unique annotation categories examined (e.g., CpG density classifiers, chromatin states, transcription factor binding, etc.)

## 3 Results

### 3.1 Participant demographics and initial findings

After performing the preprocessing procedures described in the supplementary methods, all 137 of the remaining GE measurements also had corresponding DNAm of sufficient quality in AYATS. In PREG, 131 samples had both DNAm and GE that passed quality control. While the tissue and platforms were consistent across studies, these cohorts differed in other characteristics (Table 1). PREG was an older (aged 18-40 years) and more racially diverse sample, with 49% of participants reporting African American ancestry (Supplementary Table S2). Notably, all participants in the PREG sample were pregnant women, while the AYATS sample consisted of both male (29%) and female (71%) adolescents (aged 15-20 years; Supplementary Table S1).

Genome-wide methylomic and transcriptomic data was generated using the Illumina HumanMethylation 450k BeadChip and Affymetrix HG-U133A 2.0 array, respectively. After performing quality control procedures separately in both cohorts, a differing number of probes were identified as poor quality. In AYATS, 40,392 DNAm probes and 10,809 GE probe sets were removed during preprocessing, while 63,783 DNAm and 9,473 GE measurements were removed in PREG (Table 1).

### 3.2 Associations between DNA methylation and gene expression

#### 3.3.1 Overall findings

An overview of significant DNAm-GE relationships is presented in Table 2 (see the Open Science Framework project page at https://osf.io/dk3cg/ for full lists of associations with summary statistics). Due to the differing number of measurements surviving quality control, associations with p-values < 9.68 × 10^−12^ (0.05 alpha corrected for 5,165,758,521 total tests) in PREG and p-values < 1.03 × 10^−11^ (0.05 alpha corrected for 4,857,594,560 tests) in AYATS were considered significant (Table 2). A total of 903 associations were identified in the AYATS cohort, 169 of which were in *cis* (4.72 × 10^−61^ < p < 1.01 × 10^−11^) and 734 in *trans* (2.64 × 10^−61^ < p < 1.03 × 10^−11^). Within the PREG sample, 379 DNAm-GE associations were statistically significant, of which 121 were *cis* (5.15 × 10^−58^ < p < 8.50 × 10^−12^) and 258 *trans* (2.86 × 10^−53^ < p < 9.51 × 10^−12^). Since GE probe sets measuring expression of the same gene were retained, some transcripts and CpG sites are represented more than once in the results. A total of 340 unique CpG sites and 105 unique genes comprised the 903 significant associations identified in AYATS, while 228 CpGs and 69 genes were unique in PREG across both *cis* and *trans* relationships (total n = 379). Across all categories (i.e., AYATS/PREG *cis*/*trans*), many significant relationships occurred between one transcript and one CpG site (Supplementary Figures S1 and S2), although instances in which a single CpG site was associated with multiple transcripts, and vice versa, were also common. Both positive and negative relationships were identified, although the majority of significant associations had negative coefficients (49% to 78% negative across tested categories; Table 2 and Figure 1). Effect sizes were relatively large throughout (adjusted R-squared range = 0.23-0.90), with *cis* DNAm explaining more GE variability on average (mean adjusted R-squared = 0.58 and 0.46 for AYATS and PREG, respectively) when compared to *trans* (mean adjusted R-squared = 0.48 and 0.40 for AYATS and PREG, respectively).

**Figure 1:**
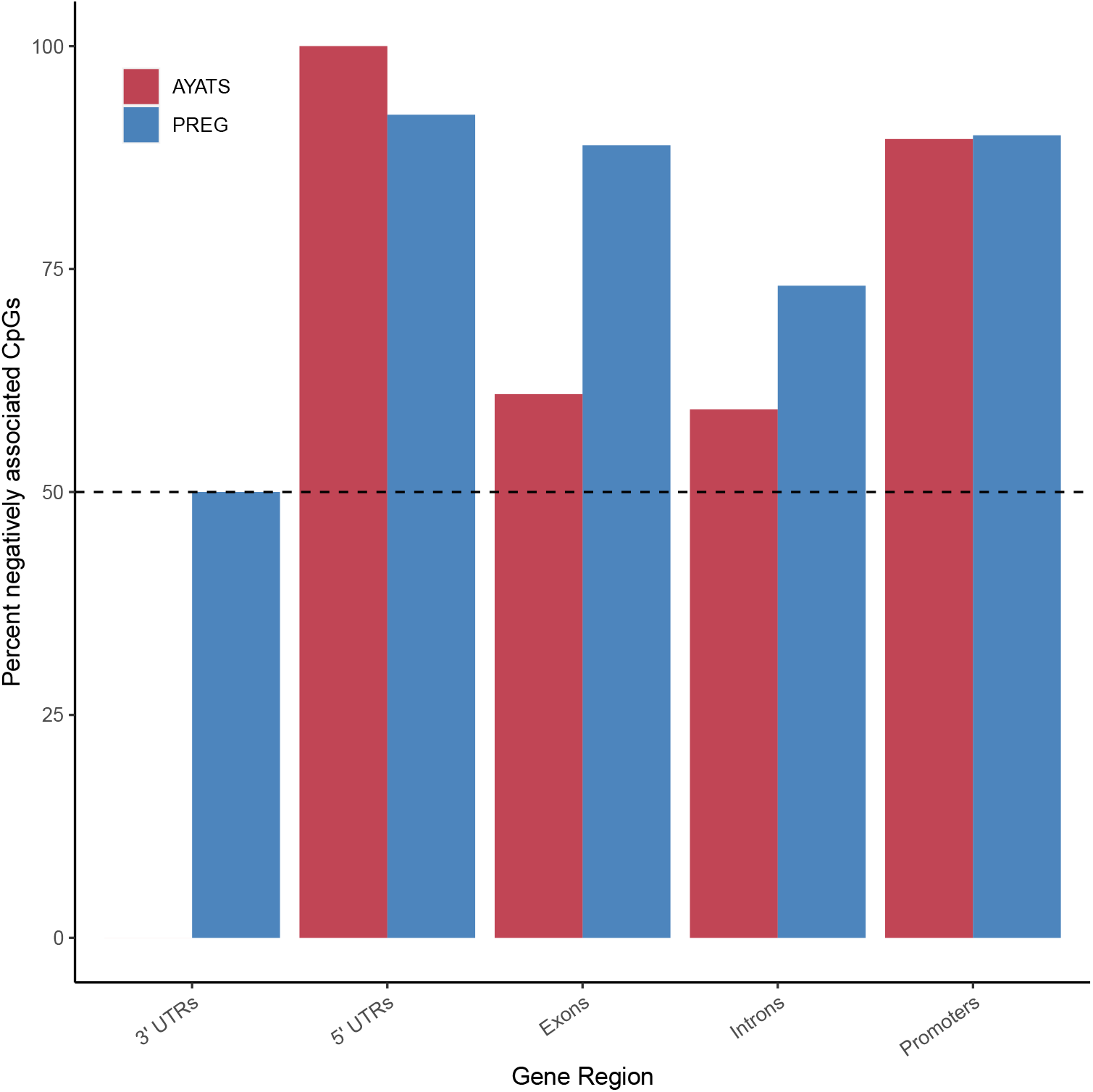
Percent negative CpG-GE associations by gene region. With the exception of 3’ untranslated regions (UTRs), the majority of *cis* CpG-GE relationships were negative across gene regions in both the AYATS (red) and PREG (blue) cohorts. Promoters and 5’ UTRs had the highest fraction of negative associations, aligning with canonical descriptions of promoter DNAm as a repressor of local gene transcription.

**Table 2:** Overview of Pairwise DNAm-GE Association Results

| Study | Significant associations<sup>1</sup> | Direction of relationship | Unique CpGs | Unique transcripts<sup>4</sup> | Mean adjusted R-squared<sup>5</sup> |
| --- | --- | --- | --- | --- | --- |
| AYATS<sup>2</sup> |  |  |  |  |  |
| *cis* | 169 | 65% negative | 107 | 42 | 0.58 (0.11) |
| *trans* | 734 | 78% negative | 266 | 96 | 0.48 (0.10) |
| PREG<sup>3</sup> |  |  |  |  |  |
| *cis* | 121 | 79% negative | 87 | 31 | 0.46 (0.10) |
| *trans* | 258 | 49% negative | 156 | 43 | 0.40 (0.08) |

<sup>1</sup> After Bonferroni adjustment for total number of tests performed within cohort
<sup>2</sup> $P\text{-val} < 1.03 \times 10^{-11}$
<sup>3</sup> $P\text{-val} < 9.68 \times 10^{-12}$
<sup>4</sup> Defined as unique by Entrez identifier
<sup>5</sup> Mean (standard deviation)
**Abbreviations.** AYATS = Adolescent and Young Adult Twin Study, PREG = Pregnancy, Race, Environment, Genes cohort.

##### AYATS Associations

The distribution of significant *cis* and *trans* connections is shown in Figure 2a. On average, each significant *cis* CpG site was associated with 1.58 transcripts (median = 1, range = 1-4; Supplementary Figure S1). *Trans* CpGs were more likely to associate with multiple transcripts than *cis* (mean = 2.75, median = 1, range = 1-22). Effect sizes, defined by adjusted R-squared values, ranged from 0.23 to 0.90. *Cis* DNAm explained more variation in GE on average (Welch’s t-test p = 2.2 × 10^−16^). DNAm-GE relationships were predominantly negative (65% of *cis* and 78% of *trans* relationships; Table 2 and Figure 1).

**Figure 2:**
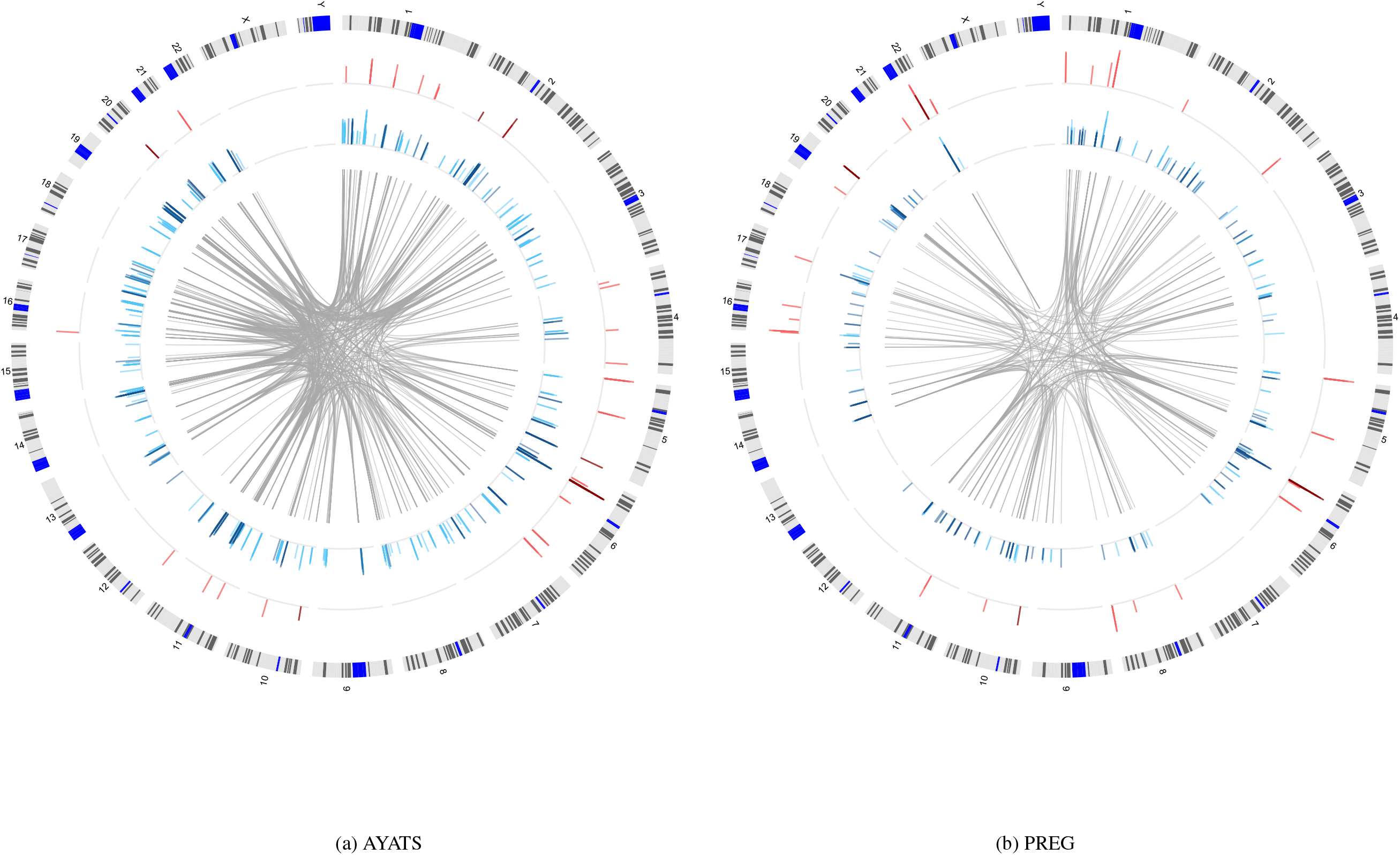
Distribution of significant connections between DNA methylation and transcript expression across the genome in the AYATS (2a) and PREG (2b) cohorts. The location of significant *cis* (red track) and *trans* relationships (blue track) across the genome (ideogram of human chromosomes, outer track) is shown. Bar graphs show the direction of the relationship (positive relationships are shown in the darker color) and the relative magnitude of the effect (height of bars; defined by adjusted R-squared values). *Trans* CpG-GE relationships often spanned chromosomes (location of associated CpG-GE pairs shown by center grey links).

##### PREG Associations

The distribution of significant *cis* and *trans* connections is shown on Figure 2b. On average, each significant *cis* CpG site associated with 1.39 transcripts (median = 1, range = 1-4; Supplementary Figure S1). Significant *trans* CpGs were more likely to associate with multiple transcripts (mean = 1.65, median = 1, range = 1-11). Effect sizes ranged from 0.31 to 0.86, with *cis* DNAm explaining more variation in GE on average (Welch’s t-test p = 5.10 × 10^−08^). *Cis* DNAm-GE relationships were predominantly negative (79%) while *trans* relationships were split almost equally between positive and negative associations (Table 2).

#### 3.3.2 Between study comparison

A total of 86 individual DNAm-transcript pairs replicated across cohorts (57 *cis* and 29 *trans*). In *cis*, 34% of significant CpG-GE pairs identified in AYATS were replicated, and 47% of those found in PREG overlapped with AYATS. In *trans*, only 4% of AYATS and 11% of PREG connections were replicated.

#### 3.2.3 Location relative to transcriptional start sites

Previous research has suggested that DNAm adjacent to the gene TSS has a stronger role in regulating proximal GE. ^24,49,59^ To explore this topic further, CpG sites associated with GE in *cis* were mapped to their associated TSS, as defined by annotations from the UCSC hg19 build knownGene track. ^60^ The location of *cis* CpG sites relative to the TSS of their associated gene is shown in Figure 3. Although many significant sites were located near the 5’ end of gene boundaries, these areas are also overrepresented in the 450k microarray. Overall, the relative proportion ([number GE-associated CpGs within 2500bp / total number GE-associated CpGs] / [number microarray CpGs within 2500bp / total number microarray CpGs]) of CpGs was higher in the GE-associated CpG sites compared to the microarray background. Interestingly, this observation was driven by CpG sites located downstream of the TSS. The relative proportion of CpG sites 2500bp upstream of the TSS was lower in GE-associated CpGs than was present on the microarray, while the opposite relationship was observed immediately downstream of the TSS (relative proportion upstream= 0.58 and 0.65; relative proportion downstream= 1.59 and 2.21 in AYATS and PREG, respectively).

**Figure 3:**
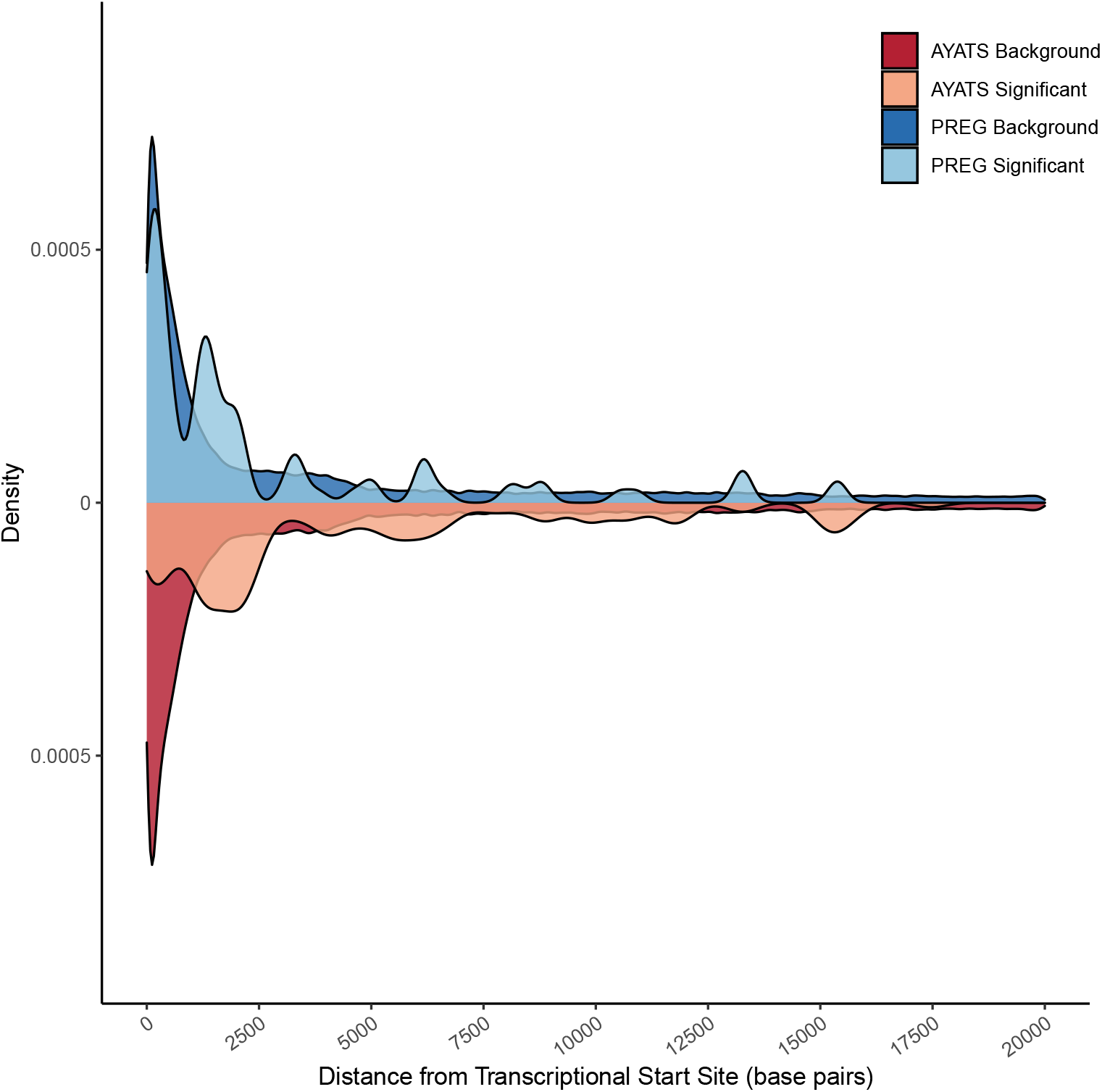
Absolute distance between CpG probes and transcriptional start sites (TSS) of proximal genes. A density plot depicting the relative distance of DNA methylation microarray probes (darker color) and significantly associated CpG sites (lighter color) from the transcriptional start sites of *cis* genes. Both the AYATS (red) and PREG (blue) cohorts showed enrichment in areas flanking active transcription. The proportion of GE-associated CpGs compared to the CpGs represented on the microarray was highest in areas directly downstream of the TSS.

### 3.3 Enrichment analyses

CpG sites were annotated by genomic regions, local CpG densities, chromatin states, bound transcription factors, and other related regulatory regions (e.g., insulator regions, regulatory RNAs). Enrichment tests were then performed within annotation type. An overview of results from enrichment analyses is outlined in Tables 3 (AYATS) and 4 (PREG). Annotation categories with p-values < 0.0025 exhibited significant depletion or enrichment, while p-values < 0.05 were considered suggestive. A number of depleted and enriched categories overlapped between the two cohorts (underlined in Tables 3 and 4). Overall, regions of high CpG density were depleted across all groups (Supplementary Table S3 and Supplementary Figures S3 - S4) while annotations indicative of regulatory activity (e.g., transcription factor binding, enhancers) were enriched among GE-associated CpGs (Tables 3 - 4).

**Table 3:** Results of AYATS Enrichment Analyses^1^

| Annotation | *cis* |  | *trans* |  |
| --- | --- | --- | --- | --- |
|  | enriched | depleted | enriched | depleted |
| Chromatin States<sup>2</sup> | **<u>TxFlnk (3),</u>**<br>**<u>ZNF/Rpts (8)</u>** | **<u>Tx (4),</u>**<br>**<u>TxWk (5),</u>**<br>Quies (15) | **<u>TssAFlnk(2),</u>**<br>**<u>TxFlnk (3),</u>**<br>**<u>TxWk (5),</u>**<br>**<u>EnhG (6),</u>**<br>**<u>Enh (7)</u>** | **<u>TssA (1),</u>**<br>**<u>Het (9),</u>**<br>TssBiv (10),<br>BivFlnk (11),<br>**<u>ReprPC (13),</u>**<br>**<u>ReprPCWk (14)</u>** |
| CpG Classifiers | **<u>South Shore</u>** | North shore,<br>**Island** | **<u>South shore,</u>**<br>**<u>Open sea</u>** | **<u>North shelf,</u>**<br>**<u>North shore,</u>**<br>**<u>Island</u>** |
| Gene Regions |  | **5' UTRs,**<br>**Promoters** | **Introns** | 5' UTRs,<br>**<u>Promoters</u>** |
| Other<sup>3</sup> | Enhancers,<br>**TF binding** | lncRNAs | **<u>Enhancers</u>** | **lncRNAs,**<br>**Insulators** |

**Abbreviations.** UTR= untranslated region, TF= transcription factor, lncRNAs= long non-coding RNAs
**Bolded items.** $P$ -val < 0.0025 (Bonferroni corrected for 20 tests)
**Underlined items.** Concordance across both the AYATS and PREG study.
<sup>1</sup> $P$ -val < 0.05
<sup>2</sup> ENCODE ChromHMM 15-state model; 1= Active transcriptional start site (TSS), 2= Flanking active TSS, 3= Flanking strong transcription, 4= Strong transcription, 5= Weak transcription, 6= Genic enhancer, 7= Active enhancer, 8= Zinc-finger genes & repeats, 9= Heterochromatin, 10= Bivalent/poised TSS, 11= Flanking bivalent TSS, 12= Bivalent Enhancers, 13= Polycomb-repressed, 14= Weak Repressed Polycomb, 15= Quiescent
<sup>3</sup> FANTOM5-defined enhancers, transcription factor binding sites derived from ENCODE TF ChIP-seq, GENCODE long non-coding RNAs

**Table 4:** Results of PREG Enrichment Analyses^1^

| Annotation | *cis* |  | *trans* |  |
| --- | --- | --- | --- | --- |
|  | enriched | depleted | enriched | depleted |
| Chromatin States<sup>2</sup> | TssA (1),<br>**<u>ZNF/Rpts (8)</u>** | **TssAFlnk (2),**<br>**Tx (4)** | **<u>TssAFlnk(2),</u>**<br>**<u>TxFlnk (3),</u>**<br>**<u>Enh (7),</u>**<br>**<u>EnhBiv (12)</u>** | **<u>TxWk (5),</u>**<br>**<u>Het (9),</u>**<br>**<u>ReprPCWk (14),</u>**<br>**<u>Quies (15)</u>** |
| CpG Classifiers | <u>South Shore</u> |  | **<u>North shore,</u>**<br>**<u>South shore</u>** | **<u>Island</u>** |
| Gene Regions |  |  | Exons | <u>Promoters,</u><br><u>3' UTRs</u> |
| Other<sup>3</sup> | **<u>TF binding</u>** |  | **<u>Enhancers,</u>**<br>**<u>TF binding</u>** |  |

**Abbreviations.** UTR= untranslated region, TF= transcription factor, lncRNAs= long non-coding RNAs
**Bolded items.** $P$ -val < 0.0025 (Bonferroni corrected for 20 tests)
Underlined items. Concordance across both the AYATS and PREG study.
<sup>1</sup> $P$ -val < 0.05
<sup>2</sup> ENCODE ChromHMM 15-state model; 1= Active transcriptional start site (TSS), 2= Flanking active TSS, 3= Flanking strong transcription, 4= Strong transcription, 5= Weak transcription, 6= Genic enhancer, 7= Active enhancer, 8= Zinc-finger genes & repeats, 9= Heterochromatin, 10= Bivalent/poised TSS, 11= Flanking bivalent TSS, 12= Bivalent Enhancers, 13= Polycomb-repressed, 14= Weak Repressed Polycomb, 15= Quiescent
<sup>3</sup> FANTOM5-defined enhancers, transcription factor binding sites derived from ENCODE TF ChIP-seq, GENCODE long non-coding RNAs

#### 3.3.1 Characterization of cis connections

South shore regions (i.e., shore regions located downstream from a CpG island) were significantly (p < 0.0025) or suggestively (p < 0.05) enriched across all groups (Figure 4). CpG islands are often associated with promoters, and both of these annotations were depleted in AYATS but were neither significantly enriched or depleted in PREG (Figures 4 and 5). Transcription factor binding sites, defined by significant peaks identified in ChIP-seq analyses of 134 transcription factors in lymphoblastoid cells, ^56^ were enriched in both cohorts (Figure 6). Chromatin state characteristics, which assign a function to genomic regions based on the presence of specific histone methylation marks, also showed some concordance between the two studies (Figure 7). Specifically, zinc-finger genes and repeats were consistently found to be enriched, whereas areas of strong transcription were consistently depleted. Regions flanking active transcription were more variably assigned, with one category found to be enriched in AYATS (areas flanking strong transcription) and another depleted in PREG (areas flanking active TSS).

**Figure 4:**
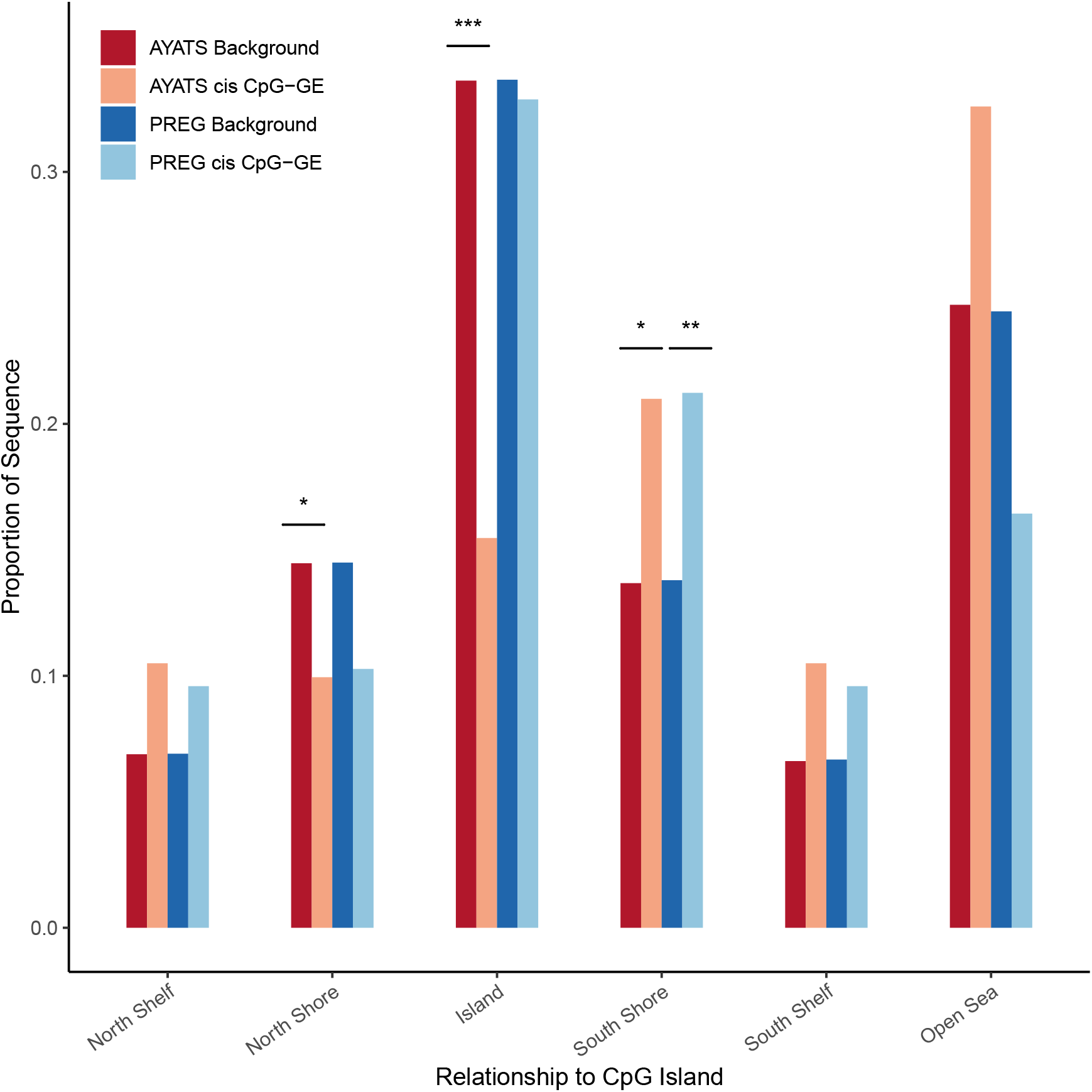
Enrichment for CpG classifiers in *cis* CpG-transcript relationships. CpG classifiers based on the distribution around CpG island regions were defined by the UCSC hg19 knownGene track. Islands and regions directly upstream from islands were depleted in AYATS. However, downstream regions bordering islands (South shores), were significantly enriched in both cohorts (*** = p < 0.0005; ** = p < 0.005; * = p < 0.05).

**Figure 5:**
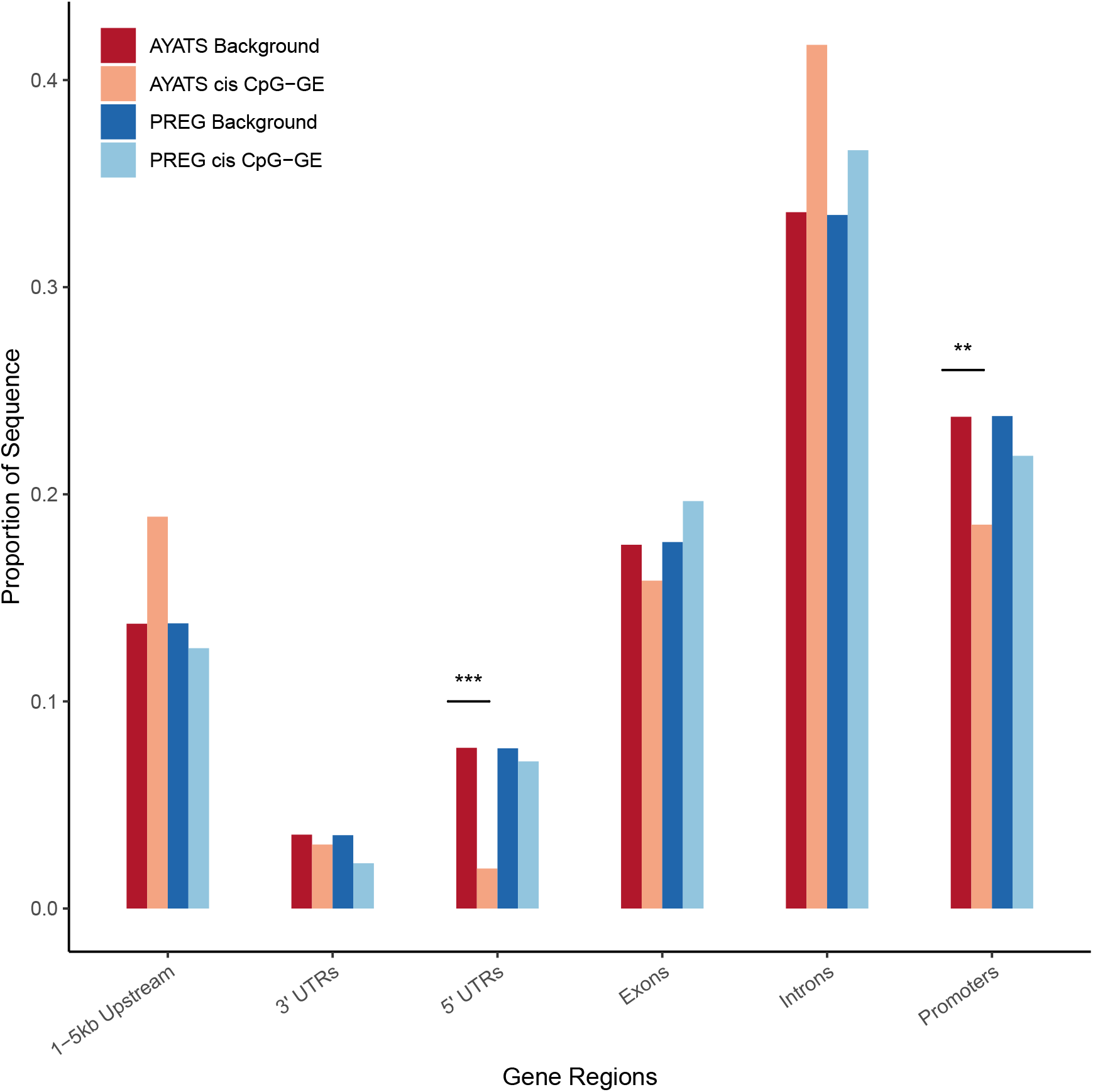
Enrichment for gene regions in *cis* CpG-transcript relationships. Gene regions were annotated based on the UCSC hg19 knownGene track. GE-associated CpG sites were depleted in 5’ untranslated regions (UTRs) and in promoters in the AYATS cohort only. (*** = p < 0.0005; ** = p < 0.005; * = p < 0.05).

**Figure 6:**
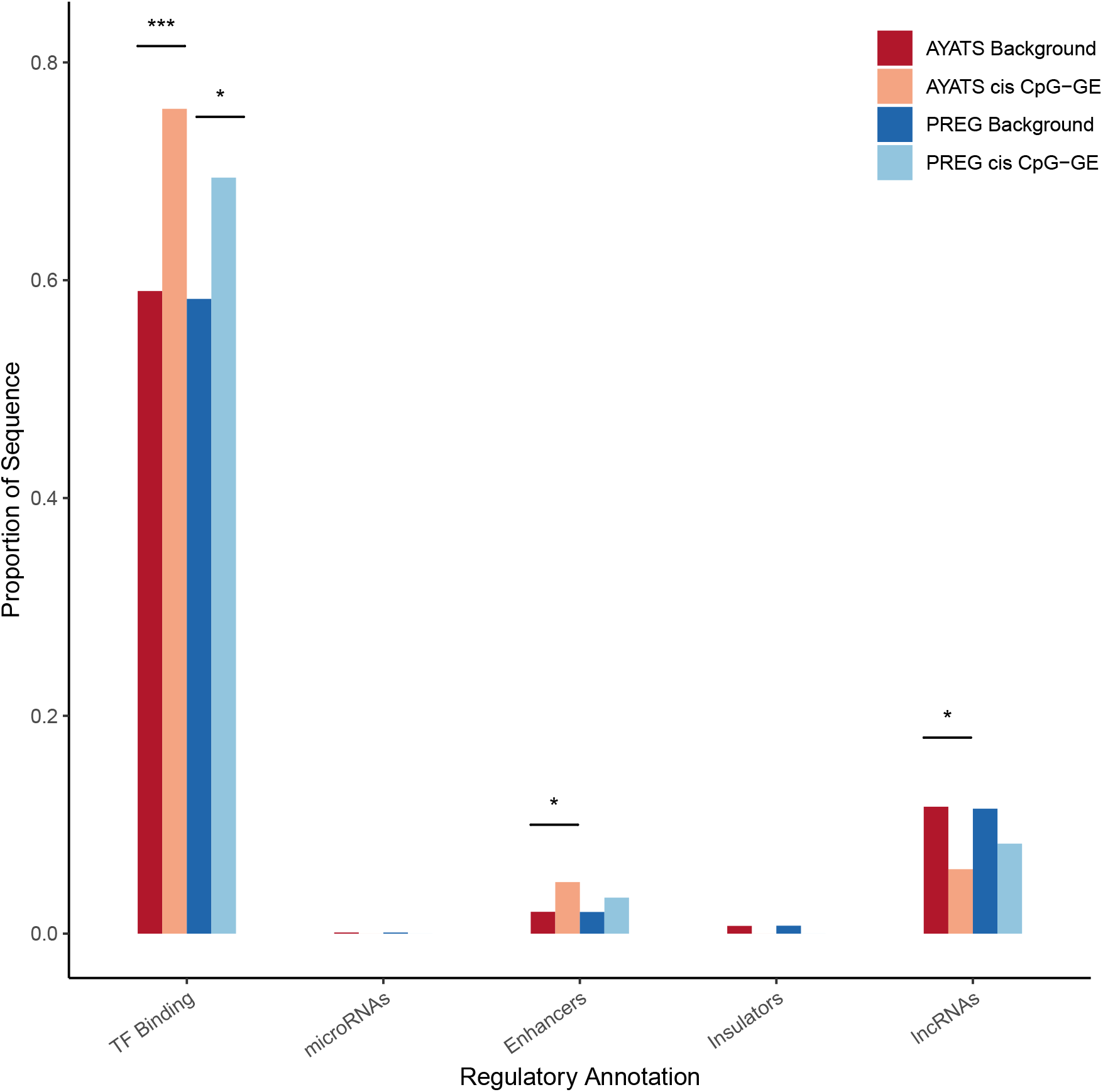
Enrichment for additional regulatory annotations in *cis* CpG-transcript relationships. Sites of transcription factor binding, as defined by ENCODE TF ChIP-seq annotations, were significantly enriched across cohorts. FANTOM5 enhancers were enriched in AYATS (*** = p < 0.0005; ** = p < 0.005; * = p < 0.05).

**Figure 7:**
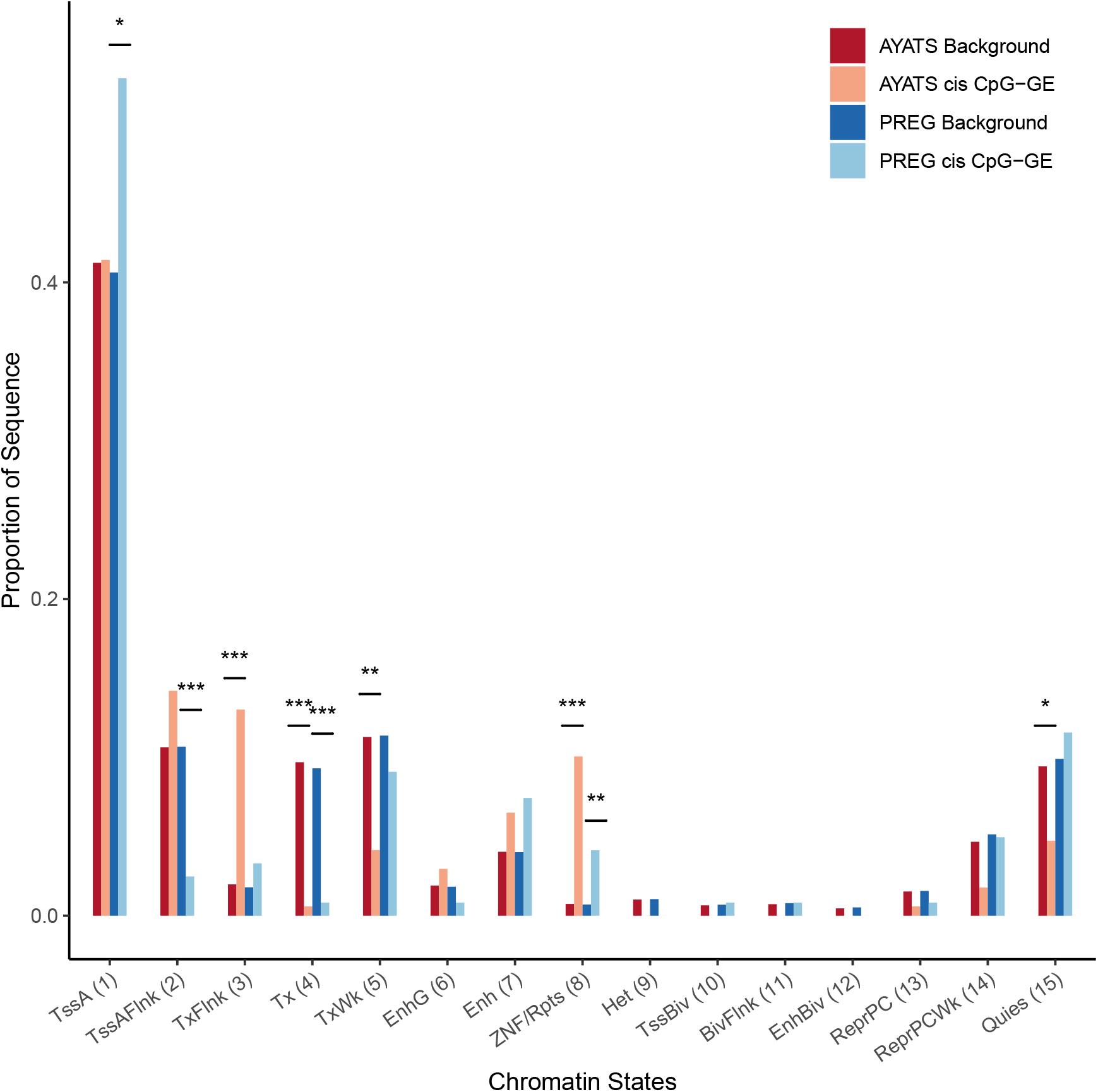
Enrichment for ENCODE chromatin states in cis CpG-transcript relationships. The 15-state ChromHMM model was used to determine regional chromatin states. Overall, GE-associated CpGs were depleted in transcriptionally active regions but enriched at zinc-finger binding sites (*** = p < 0.0005; ** = p < 0.005; * = p < 0.05). **Abbreviations:** 1= Active transcriptional start site (TSS), 2= Flanking active TSS, 3= Flanking strong transcription, 4= Strong transcription, 5= Weak transcription, 6= Genic enhancer, 7= Active enhancer, 8= Zinc-finger genes & repeats, 9= Heterochromatin, 10= Bivalent/poised TSS, 11= Flanking bivalent TSS, 12= Bivalent Enhancers, 13= Polycomb-repressed, 14= Weak Repressed Polycomb, 15= Quiescent

#### 3.3.2 Characterization of trans connections

Like *cis* CpG-GE pairings, CpGs associated with GE in *trans* were overall depleted in areas of high CpG density (i.e., within CpG islands) and within the promoter regions of genes, while South shore regions were enriched (Tables 3 - 4 and Figures 8 - 9). Again, GE-associated CpG sites were overrepresented in areas of known regulatory importance, such as sites of transcription factor binding and enhancer regions (Figure 10). With the exception of lncRNAs, which were depleted in AYATS, noncoding RNAs were neither over- or underrepresented. The chromatin state analysis highlighted distinct differences between *cis* and *trans* results. Chromatin states reflecting enhancer regions were enriched in both AYATS and PREG, as were areas flanking sites of active transcription. Repressed states, including heterochromatic regions and polycomb-repressed regions, were consistently depleted (Figure 11).

**Figure 8:**
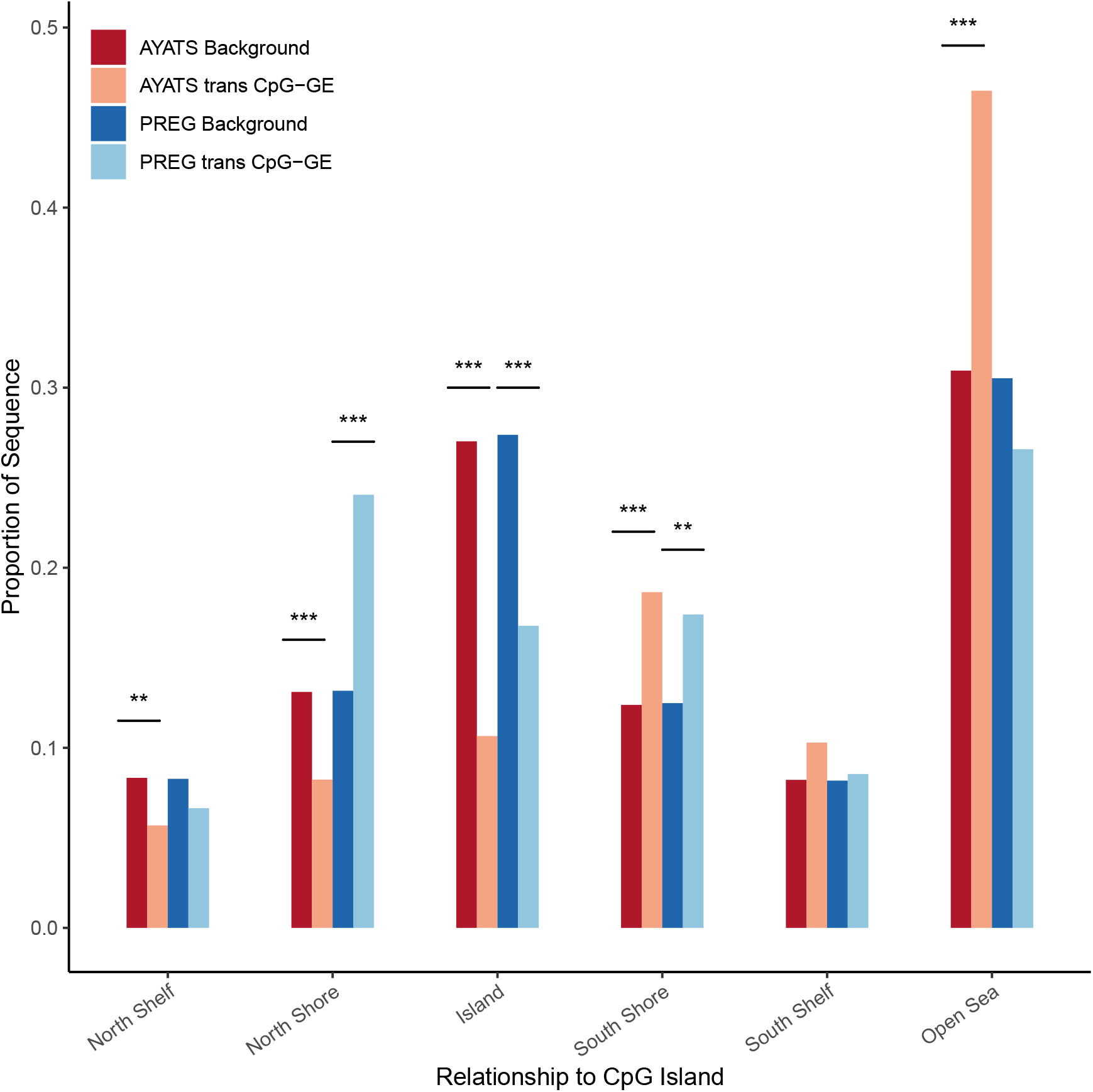
Enrichment for CpG classifiers in *trans* CpG-transcript relationships. CpG classifiers based on the distribution around CpG island regions were defined by the UCSC hg19 knownGene track. Islands were depleted while downstream regions bordering islands were significantly enriched in both cohorts. The North shore region directly upstream of CpG islands was more variable, with significant CpGs showing depletion in AYATS and enrichment in PREG (*** = p < 0.0005; ** = p < 0.005; * = p < 0.05).

**Figure 9:**
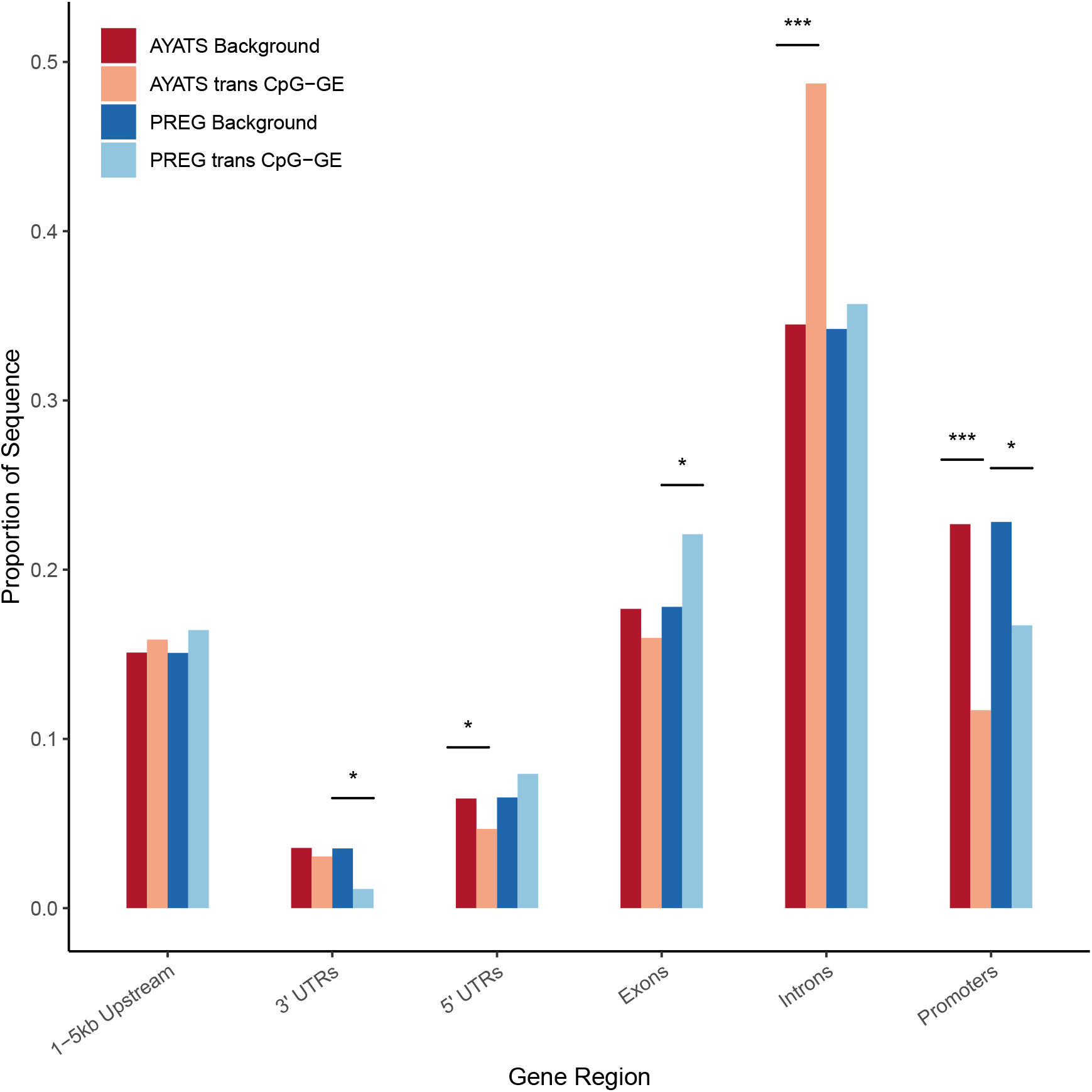
Enrichment for gene regions in *trans* CpG-transcript relationships. Gene regions were annotated based on the UCSC hg19 knownGene track. GE-associated CpG sites were depleted in 3’ untranslated regions (PREG), 5’ untranslated regions (AYATS), and in promoters (AYATS and PREG). Exons and introns were enriched in PREG and AYATS, respectively (*** = p < 0.0005; ** = p < 0.005; * = p < 0.05).

**Figure 10:**
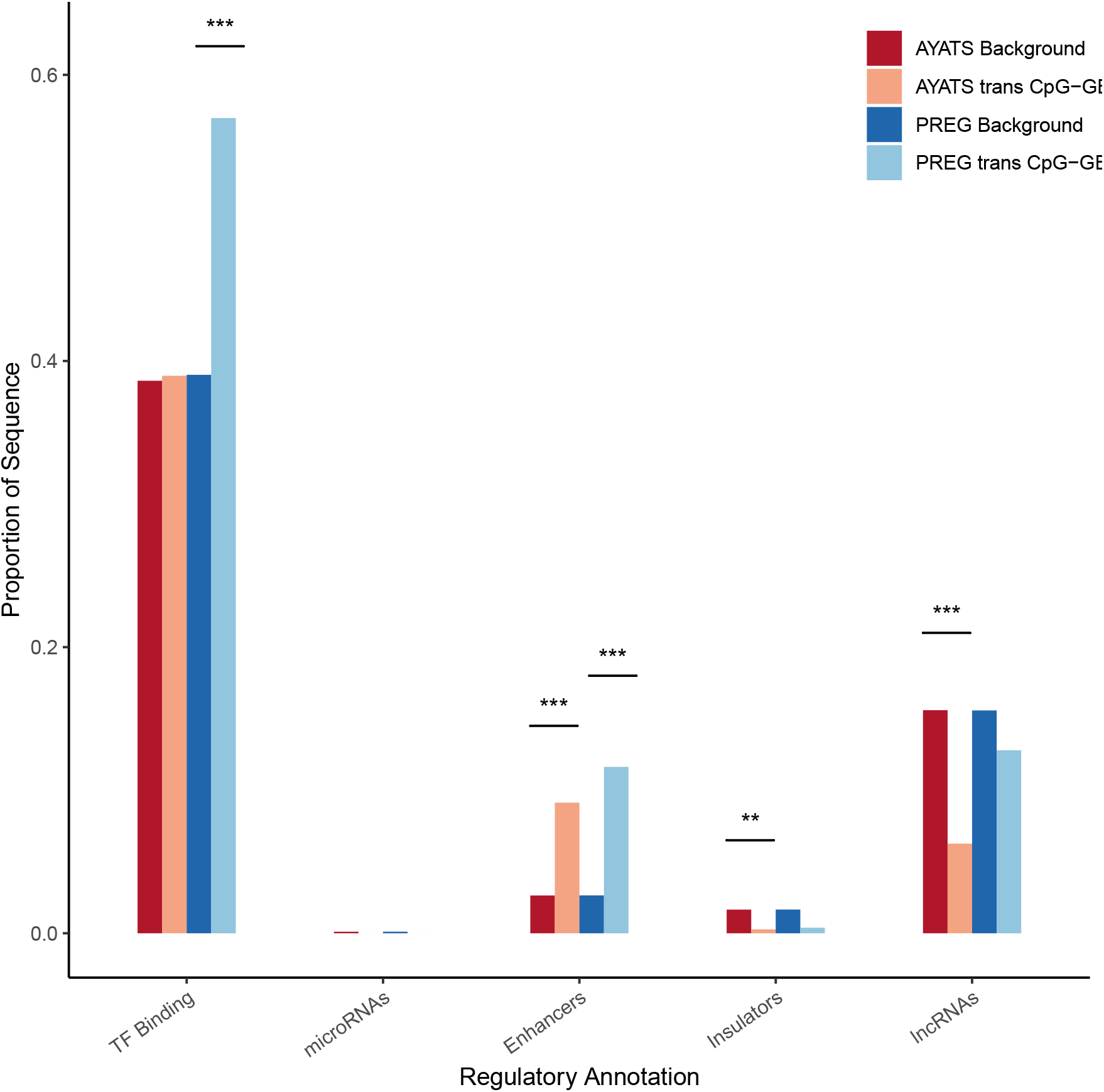
Enrichment for additional regulatory annotations in *trans* CpG-transcript relationships. Sites of transcription factor binding, as determined by ENCODE TF ChIP-seq, were significantly enriched in the PREG cohort. Enhancers were enriched across cohorts, and insulator regions were depleted in AYATS (*** = p < 0.0005; ** = p < 0.005; * = p < 0.05).

**Figure 11:**
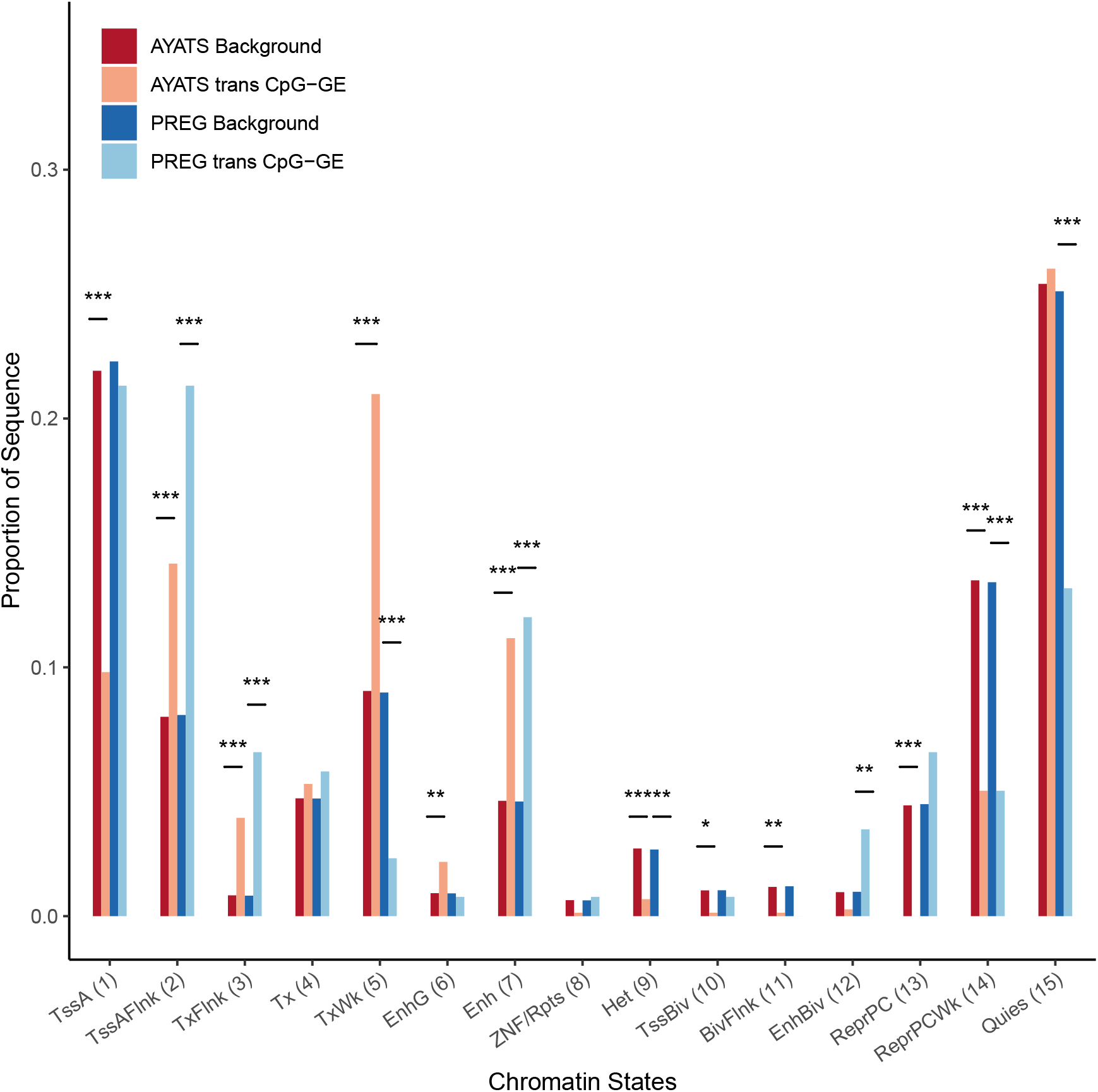
Enrichment for ENCODE chromatin states in *trans* CpG-transcript relationships. The 15-state ChromHMM model was used to determine regional chromatin states. Overall, GE-associated CpGs were depleted in repressive states but enriched at enhancers and areas flanking actively transcribed genes (*** = p < 0.0005; ** = p < 0.005; * = p < 0.05). **Abbreviations:** 1= Active transcriptional start site (TSS), 2= Flanking active TSS, 3= Flanking strong transcription, 4= Strong transcription, 5= Weak transcription, 6= Genic enhancer, 7= Active enhancer, 8= Zinc-finger genes & repeats, 9= Heterochromatin, 10= Bivalent/poised TSS, 11= Flanking bivalent TSS, 12= Bivalent Enhancers, 13= Polycomb-repressed, 14= Weak Repressed Polycomb, 15= Quiescent

#### 3.3.3 Functional enrichment analysis

The Gene Ontology (GO) Consortium and the Kyoto Encyclopedia of Genes and Genomes (KEGG) were used to assess overrepresented gene functions and pathways within significant *cis* results (see supplement for more information). Terms with a false discovery rate (FDR) < 0.05 were deemed significant. ^61^ Common themes were uncovered in both cohorts (Supplementary Tables S4 - S11), and include functions related to the activation and regulation of immune response and cellular detoxification. A total of 33 significantly enriched GO terms overlapped between the two cohorts, and all significant KEGG pathways identified in AYATS (n = 31) were also found in PREG (n = 39). However, these consistences were supported by relatively few genes.

## 4 Discussion

Although genome-wide epigenetic studies aim to uncover the role of DNAm in disease development and progression, they often do not utilize an experimental framework that provides evidence for a mechanistic relationship. Most EWAS operate under the assumption that DNAm influences proximal gene transcription. However, the absence of measured GE makes relying on this interpretation difficult, especially as mounting evidence suggests that DNAm does not always follow a canonical *cis* relationship. ^15,16^ Given the complicated network of interactions between DNAm, GE, higher-order chromatin modifiers, and other regulatory elements, it is challenging to draw accurate conclusions about the downstream functional effects of altered DNAm without, at minimum, integrating concomitant measurements of GE. ^15,16,20,56^ To investigate the relationship between DNAm and GE further, this study tested genome-wide associations between DNAm and GE in peripheral blood collected from two cohorts. Both canonical and non-canonical relationships were identified, highlighting potential inaccuracies in the current functional interpretation of trait-associated CpG sites within the frequently adopted EWAS framework.

Across cohorts, DNAm was significantly associated with both proximal (*cis*) and distal (*trans*) GE. The primary findings of this study align with other reports of long-range DNAm-GE relationships, adding to the growing body of literature questioning the accuracy of current EWAS interpretations. ^17,24,62^ The effect sizes detected among DNAm-GE pairs were relatively large, with DNAm predicting 42-50% percent of GE variability on average. Although this result is likely influenced by a lack of statistical power to detect more attenuated relationships, it reiterates that while DNAm may not be an appropriate proxy for changes in GE, strong links between the two measurements exist. Approximately 23% of connections identified in PREG were also significant in the AYATS cohort, suggesting a consistent program of gene regulation even among the disparate cohorts tested. While this proportion is similar to DNAm-GE connections identified in peripheral blood and isolated monocytes, ^24^ discrepancies between the two cohorts could be related to differences in statistical power or differences in demographic and clinical features (i.e., genetic ancestry, developmental stage, etc.). On average, *cis* connections were more likely to replicate between studies and account for larger proportion of GE variability when compared to within-cohort *trans* associations. Larger samples are likely necessary to detect more subtle *cis* and *trans* CpG-GE pairings and provide a balanced assessment of the expected replication across samples.

Interpreting DNAm-disease relationships is hindered not only by limitations in identifying DNAm-GE pairs, but also by challenges in predicting the precise functional impact of altered DNAm on an associated gene’s expression. Both negative and positive relationships between *cis* and *trans* DNAm-GE pairs were identified (Figure 1). Although DNAm is usually considered a repressive mark, inhibiting GE by either blocking transcription factor binding or by promoting a more condensed DNA conformation, ^16^ positive DNAm-GE relationships could be explained by several mechanisms. Within genomic regulatory elements, transcription factors with repressive, rather than activating properties, may bind unmethylated sequences. ^23^ Furthermore, many transcription factors actually exhibit an increased affinity for heavily methylated sites. ^21^ Besides influencing the binding affinity of regulatory proteins, DNAm patterns may also reflect a history of transcription factor binding, a phenomenon that cannot be separately identified by a classic EWAS design. ^63^ In recent years, speculation has emerged regarding potential alternative roles of DNAm in the cell, including theories that DNAm may serve to direct splicing regulation or in maintaining genomic stability within specific regions. ^5,64–67^ Although this study found that the associations were predominantly negative across the majority of gene regions (Figure 1), these findings agree with other reports that strong positive DNAm-GE relationships exist. ^24,25,49^

Given the large effect size distribution of detected associations and the modest number of participants in each cohort, it was expected that only a small proportion of DNAm-GE connections would reach statistical significance. Instead of only focusing on individual connections, this study sought to outline genome-wide trends by identifying attributes of GE-associated CpG sites. Despite the modest number of DNAm-GE pairs overlapping across cohorts, GE-associated CpG sites displayed similar annotation characteristics (Tables 3 and 4). Annotations uniquely characterized attributes of *cis* and *trans* GE-associated CpGs, indicating that separate paradigms may exist for proximal and distal connections. In general, DNAm within intermediate CpG density regions were more likely to be associated with GE. Regions of intermediate CG density are more variable compared with low- or high-density regions, and appear more dynamic across tissues and developmental stages. ^68,69^ Conversely, CpG sites in high density regions, which are most often associated with CpG islands and promoter regions, were consistently depleted. Interestingly, both transcription factor binding and chromatin states in regions near active TSS were enriched, with those regions directly downstream of the TSS particularly characterized by a high proportion of GE-associated CpGs (Figure 3). Other studies have noted a similar relationship with DNAm located in the first intron, while also observing high transcription factor activity typical of intronic enhancers within these areas. ^59,70,71^ Although mechanisms of DNAm transcriptional inactivation usually focus on the hypermethylation of CpG sites within promoter and island regions, these results agree with other studies showing enrichment for "off-island" DNAm among GE-associated CpG sites. ^24,68^

The traditional DNAm regulatory paradigm suggests the importance of DNAm in promoter regions, but results from this study instead reiterate the significance of DNAm within enhancer regions. ^24,27^ Multiple enhancer definitions (i.e., enhancer-like chromatin states and enhancer annotations generated from cap analysis of gene expression [CAGE]) were enriched within *trans* results. Enhancers have an established role in long-range gene regulation, ^9,72^ often looping over more proximal genes to interact with those farther away. ^73^ Enhancer regions are often characterized by intermediate DNAm and chromatin accessibility, demonstrate greater DNAm variability than promoters, ^52,68^ and exhibit ongoing *de novo* methylation and demethylation activity. ^48^ The high rate of DNAm remodeling within enhancers, coupled with the strong DNAm-GE relationships found within these regions, align with hypotheses that suggest environmental exposures can influence complex disease risk through epigenetic mechanisms of transcriptional dysregulation. While mapping enhancers to their putative genes is a fundamental aim in identifying transcriptional regulatory networks, current methods are still under development, ^73^ adding to the uncertainty in predicting the downstream functional effects of DNAm within these distal regulatory regions. Further challenges arise from evidence that many genes actually interact with multiple enhancers, and that these compounded interactions can result in additive effects on target GE. ^73^ However, the relationships between DNAm outside of proximal regulatory elements and GE again question the generalizability of the canonical DNAm regulatory mechanism and suggests that EWAS should transition away from relying on this paradigm to interpret underlying disease biology.

This study serves to improve understanding of the relationships between DNAm and GE across the genome and adds to the growing body of literature which cautions against misconstruing modified DNAm as changes in proximal GE. Overall, these results highlight issues with restricting DNAm-transcript annotations to small genomic intervals and question the validity of assuming a canonical *cis* DNAm-GE pathway when investigating epigenetic mechanisms. The results from this study underscore concerns in predicting the biological mechanisms underlying disease from DNAm measurements alone. EWAS relying on a canonical DNAm-mediated transcriptional regulatory mechanism to interpret DNAm-trait associations may reach inaccurate conclusions about disease pathoetiology. Even modified EWAS that incorporate GE information by performing a transcriptome-wide association study alongside testing for DNAm-disease associations should interpret findings with care, since in this study design *a priori* assumptions link CpG sites to putative genes and DNAm-GE relationships remain uninvestigated.

Based on these results, epigenetic research should continue moving towards multi-omic approaches that integrate DNAm with other levels of data (e.g., GE, genotypes, transcription factor binding) to study complex traits. Although DNAm-GE relationships are highly complex, the integration of DNAm with data outlining regional chromatin architecture and transcription factor activity may assist in predicting the functional impact of altered DNAm. ^24,74^ However, as an emerging and heterogeneous field, several obstacles can interfere with the implementation and interpretation of multi-omic studies. ^75–78^ Standardized analytical pipelines have yet to be developed, leading to difficulties in cross-study comparisons and in assessing rigor. ^77^ Currently, only a handful of studies have tested genome-wide associations between GE and DNAm, ^24,62,68,79,80^ but variability in the methodologies used has lead to difficulties in determining the replicability and generalizability of identified relationships. Although cross-study comparisons are challenging, several consistent themes have emerged from this modest body of literature. This study replicates the overrepresentation of GE-associated CpGs within enhancers and at transcription factor binding sites, as well as the depletion within islands and promoter regions. ^24^ Moving forward, continued examination of DNAm-GE relationships in large, diverse cohorts should be prioritized to advance our understanding of the role of DNAm within the cell and disease biology.

## 5 Strengths and Limitations

To our knowledge, this was the first study to assess the global relationship between peripheral blood DNAm and GE in both a primary and replication sample. However, results of this study should be considered in the context of the following limitations. First, both DNAm and GE were measured by microarray technologies that provided coverage within well-characterized locations, but were unable to assay the full extent of RNA and CpG sites in the genome. ^81^ Second, only relationships of relatively large effect size were detected in this study (adjusted R-squared range = 0.23-0.90). Especially given that a conservative multiple testing correction was applied, it is assumed that many more DNAm-GE connections exist but were undetected in this study, which could influence the results of the feature enrichment tests. Third, both cohorts were analyzed cross-sectionally, a study design that is unable to provide evidence for causation or directionality. ^15,82^ Mechanisms of reverse causation, in which changes to DNAm occur in response to modified GE, have been observed. ^83^ Therefore, it is unknown whether changes to DNAm are actually proceeding changes in GE as described in the canonical mechanism. Fourth, some annotations were derived from experiments conducted on a well-described lymphoblastoid cell line (GM12787), which was selected based on data that supports the genetic and functional similarity to mature blood cells (i.e., T cells and B cells). ^84^ One benefit of using this approach is that annotations were kept consistent across the different functional enrichment categories (e.g., chromatin landscapes, enhancer definitions, etc.). It remains important to consider that this study focused on the association between DNAm at individual CpG sites and GE. In actuality, regional changes in DNAm may also be co-regulating GE together. ^62,85^ While correlations between proximal GE-associated CpG sites did not suggest a predictable method in which to aggregate measured sites, future studies can examine how regions of CpGs work in concert to regulate GE. ^62^ Finally, this study only investigated DNAm and GE in the peripheral blood and may not generalize to other tissues. ^59^ Future analyses with more comprehensive measurements in alternative tissues will be crucial for characterizing genome-wide trends across cell types.

## 6 Additional information

### 6.1 Acknowledgements

The PREG study was supported by the NIMHD (P60MD002256, PI: York) and by the Clinical and Translational Science Award (CTSA) award No. UL1TR000058 from the National Center for Advancing Translational Sciences. Support for the AYATS study was received by a NIH/NIMH R01 (MH101518), R21 (MH106924), and a NARSAD Independent Investigator Award from the Brain and Behavior Research Foundation (PI: Roberson-Nay).

### 6.2 Data sharing

Sharing PREG and AYATS study data is limited by Institutional Review Board agreements and participant consent forms, which restrict openly sharing individual-level measures. Anyone interested in data access or collaboration is encouraged to contact Dr. Timothy P. York or Dr. Roxann Roberson-Nay for more information.

### 6.3 Author contributions

All authors assisted with the design of the study. EL cleaned the data, performed the analyses, and wrote the initial draft of the manuscript. RRN, VV, BR, JL and TY provided substantive feedback and revisions, and TY and RRN planned and secured funding for the PREG and AYATS studies.

## 6.5 Abbreviations

AYATS: Adolescent and Young Adult Twin Study
DMPs: Differentially methylated positions
DNAm: DNA methylation
EWAS: Epigenome-wide association study
FDR: False discovery rate
GE: Gene expression
GO: Gene ontology
KEGG: Kyoto Encyclopedia of Genes and Genomes
PREG: Pregnancy, Race, Environment, Genes study
TSS: Transcriptional start site
UTRs: Untranslated regions

