## Supplemental Materials for "Large-scale integration of DNA methylation and gene expression array platforms"

### 1 Demographic Information

#### 1.1 AYATS

Participants were primarily recruited through the population-based Mid-Atlantic Twin Registry (MATR).[1,2] The original Research Domain Criteria (RDoC) sample did not select for any particular psychiatric disorder, but a subset of twin pairs discordant and concordant for a lifetime history of MD were preferentially chosen for the DNAm and GE analyses. A total of 83 monozygotic twin pairs were selected for their adherence to the study's inclusion criteria and no exclusion criteria.[1,3] Inclusion criteria at enrollment included twin monozygosity, aged between 15-20 years, and living primarily at home. Exclusion criteria were the current use of psychotropic medications or those with psychotropic effects, diagnoses of autism spectrum disorder or intellectual disability, prior testing indicating an IQ below 70, idiopathic seizure, current or past episodes of psychosis, any serious unstabilized illness, low production of human growth hormone, sensory integration disorder, congenital adrenal hyperplasia, adrenal inefficiency, deaf with bicochlear implants, current or previous cancer diagnosis, and pregnancy. All twins self-identified as Caucasian or Latinx. Peripheral blood collected from participants at a single time point was assayed for both DNAm and GE. A total of 137 participants (65 twin pairs plus 7 singletons) had both DNAm and GE measurements that passed quality control procedures. An overview of sample characteristics is provided in Table S1 and further demographic information can be accessed at Roberson-Nay et al, 2018.[3]

#### 1.2 PREG

Pregnant women were recruited from 2013-2016 at the Virginia Commonwealth University Medical Center in Richmond, Virginia.[4] Of the 240 women who were initially enrolled in the study, 177 met all birth and pregnancy inclusion criteria and no exclusion criteria. Eligibility criteria at enrollment included singleton pregnancy, enrollment prior to 24 weeks gestation, maternal age 18-40 years old at enrollment, no diagnosis of diabetes, and no use of reproductive technology to conceive the current pregnancy. Exclusion criteria at birth included either infant or placental abnormalities, polyhydramnios/oligohydramnios, preeclampsia/pregnancy-induced hypertension (PIH)/haemolysis, elevated liver enzymes, low platelet count (HELLP), Rh sensitization, the use of cervical cerclage, medically necessitated preterm delivery, drug abuse, and fewer than 2 time points of data. Peripheral blood samples were collected up to four times throughout pregnancy. DNAm was assessed at all time points whereas GE was measured once at the final collection during weeks 37–42 of gestation. A total of 159 women had concomitant DNAm and GE measured. After removing poor quality samples defined by the procedures in Section 3, 131 participants had concurrent DNAm and GE measurements. An overview of sample characteristics is provided in Table S2 and further study information can be accessed at Lapato et al., 2018.[4]

Table S1: AYATS cohort characteristics

| | MD Cases | Controls |
|---|---|---|
| N | 35 (26%) | 102 (74%) |
| Age (years) | 17.11 (1.43) | 16.9 (1.22) |
| Ethnicity (Caucasian)[1] | 34 (97.1%) | 98 (96.1%) |
| Sex (Female) | 25 (71.4%) | 72 (70.6%) |
| Nicotine (Current smoking status) | 3 (8.6%) | 2 (2%) |
| Phenotypic characteristics | | |
| Psychotropic medication history | 4 (11.4%) | 2 (2%) |
| SMFQ | 8.83 (6.0) | 4.00 (3.4) |
| 1 Depressive Episode | 16 (45.7%) | - |
| 2-3 Depressive Episodes | 15 (42.9%) | - |
| 4-5 Depressive Episodes | 4 (11.4%) | - |
| Age of onset (years) | 14.56 (3.07) | - |

Includes all participants with DNA methylation and concomitant gene expression data after quality control procedures.
M (SD) or N (%)
[1] All participants identified as either Caucasian or Latinx
**Abbreviations.** MD= major depression, AYATS= Adolescent and Young Adult Twin Study, SMFQ= Short Mood and Feelings Questionnaire.

Table S2: PREG cohort characteristics

| | Participants |
|---|---|
| N | 131 |
| Ethnicity (Caucasian)[1] | 67 (51.1%) |
| Age | 29.1 (5.0) |
| Sex (Female) | 131 (100.0%) |
| Nicotine (Current smoking status)[1] | 6 (4.6%) |
| Phenotypic characteristics | |
| GA Delivery | 277.4 (8.1) |
| Primiparous | 80 (61.1%) |
| Prenatal vitamin[2] | 82 (62.6%) |

Includes all participants with DNA methylation and concomitant gene expression data after quality control procedures.
M (SD) or N (%)
[1] All participants identified as either Caucasian or Black
[2] Assessed at first study visit
**Abbreviations.** GA= gestational age at delivery (in days).

#### 2 Platform descriptions

Both PREG and AYATS utilized the same platforms to assay DNAm and GE. DNAm was measured using the Illumina 450k HumanMethylation BeadChip, while GE measurements were generated by the Affymetrix GeneChip Human Genome U133A 2.0 microarray.

**Illumina HumanMethylation 450k BeadChip.** The Illumina 450k microarray measures DNAm at 485,512 CpG sites across the genome. Almost all RefSeq genes (99%) and 96% of CpG islands are represented by at least one CpG site on the array. Although more intergenic CpG sites were added since the previous iteration, the Illumina HumanMethylation27 array, promoter CpG sites are still the most well-represented gene region (41%).[5]

**Affymetrix HG-U133A 2.0.** The Affymetrix U133A microarray manufacturers focused on representing well-characterized genes and selected sequences from GenBank, dbEST, and RefSeq. The array contains 21,722 probe sets which measure the expression of 14,564 genes (as defined by UniGene clusters and subclusters, respectively).[6]

#### 3 Molecular measurements and raw data processing

All data processing steps were performed separately for each cohort using Bioconductor packages in the R programming environment.[7]

##### 3.1 DNA methylation measurement

DNAm was measured once per participant in AYATS but at multiple time points in the PREG cohort. The DNAm measurement concomitant with GE was selected to match the AYATS study design and facilitate cross-study comparisons. Otherwise, preprocessing methods were consistent across the two studies.

Steps to generate raw measures of DNAm by microarray typically involve collecting tissue samples, extracting the DNA from those cells, performing sodium bisulfite conversion, hybridizing to the array, and reading the proportion of methylated to unmethylated signals. Whole blood was first collected into EDTA tubes and genomic DNA was isolated from 10 mL whole blood according to standard methods using the Puregene DNA Isolation Kit (Qiagen; Valencia, CA). An aliquot of 1 $\mu$g DNA per participant was sent for bisulfite conversion (HudsonAlpha Institute for Biotechnology; Zymo Research EZ Methylation Kit). Genome-wide methylation was assayed according to the manufacturer's protocol at the HudsonAlpha Institute for Biotechnology (Illumina, San Diego, CA, USA). The 450k array consists of 6 rows and 2 columns that can hold up to twelve samples per array. Samples were randomized to minimize any potential batch effects that might arise from this structure (i.e., differences by array, row, or column).

##### 3.2 DNA methylation preprocessing

Raw Intensity Data files (*.idat) files were generated after scanning the microarray chips and read into R for processing using the `minfi` package from the Bioconductor suite.[8] Standard quality control pipelines for processing the Illumina 450k array were followed.[9–13] Briefly, poor quality probes and samples were identified and data was normalized before other potentially problematic probes are filtered. High detection p-values often indicate technological issues with the scanning of arrays (e.g., spatial artifacts). Probes with detection p-values greater than 0.01 were considered poor, and any probes that failed in more than 1% of samples were removed. Poor quality samples were defined as having both methylated and unmethylated median signal intensities less than 10.5 units and removed. Visual inspection confirmed that outlier samples had been eliminated. Quantile normalization was implemented to adjust for background correction, color bias, and probe type bias.[8,14]

In addition to identifying those poorly performing probes, several other potentially confounding probes were removed. Cross-hybridizing probes, probes targeting sites on the X and Y chromosomes, and those probes with polymorphisms at the CpG site and single-base extension were removed prior to analysis.[15] Measurements from sex chromosomes were removed in PREG and AYATS to maintain a reasonable consistency in methodology across the two studies and facilitate any potential cross-study comparisons.

In order to adhere to assumptions of normality in testing, the ratio of methylated signal to total signal (i.e., Beta values) were converted to M-values, or the log2 ratio of the intensities of methylated probe versus unmethylated probe.[16] Finally, M-values were further adjusted for slide-related technical artifacts prior to analysis using an empirical Bayes framework.[17]

#### 3.3 Gene expression measurement

Gene expression of more than 14,500 well-characterized genes was measured on the Affymetrix Human Genome U133A 2.0 Array.[6] Blood was collected in EDTA tubes and processed immediately in the PREG cohort, whereas AYATS relied on frozen blood samples collected in PAXgene RNA tubes. Quality of the initial RNA and synthesis products were monitored closely throughout procedures.[18] First, RNA was extracted from 10 ml of whole blood and RNA purity was judged by spectrophotometry at 260, 270, and 280 nm. Following the standard Affymetrix protocol, initial RNA served as a template for cDNA synthesis, from which biotinylated cRNA is then synthesized. RNA integrity as well as cDNA and cRNA synthesis products were assessed by running 1 $\mu$L of every sample in RNA 6000 Nano LabChips on the 2100 Bioanalyzer (Agilent Technologies, Foster City, CA). Biotinylated cRNA was generated with the GeneChip 3'IVT Express kit and 20 $\mu$g of the cRNA product were fragmented and 15 mg of the fragmented product were hybridized for 18 to 20 hours into Affymetrix HG-U133A 2.0 microarrays. Each microarray was washed and stained with streptavidin-phycoerythrin and scanned at a 6 mm resolution by the Agilent G2500A Technologies Gene Array scanner (Agilent Technologies, Palo Alto, CA) according to the GeneChip Expression Analysis Technical Manual procedures (Affymetrix, Santa Clara, CA).[18]

#### 3.4 Gene expression preprocessing

Raw data CEL files were processed using the `affy` package from the Bioconductor suite.[19] The overall quality of each array was assessed by monitoring the 3'/5' ratios for 2 housekeeping genes (GAPDH and Beta-actin), the percentage of 'present' genes, and the relative log expression (RLE) which assesses the presence of unwanted variation.[20] These parameters plus several others (e.g., spatial artifacts that identify damaged arrays) were assessed visually.[21] Correlations between technical variables (e.g., cell types, hybrization date, scan date) and principal components assessed for the presence of batch effects or the influence of other potential confounders.[22] Samples identified as outliers on multiple quality control procedures were removed.

The recommended adjustments to raw expression intensities were applied prior to analyzing.[23] Background correction, log transformation, quantile normalization, and estimation of probe set expression summaries was performed using the robust multiarray analysis (RMA) within the `affy` Bioconductor package.[24] Finally, expression values were adjusted for scan group using an empirical Bayesian framework method present in the `sva` R package.[25]

Summarized probe sets were annotated using data derived from the University of California Santa Cruz (UCSC) Known Genes database, and linked to sequence position on the GRCh37 (hg19) assembly using Entrez identifiers.[26,27] Several probe sets are removed during processing steps to ensure that only the expressed genes of good quality are retained for analyses. Some redundancy is present on the array, so that 37% of genes are measured by more than one probe set.[28] For example, although 10,913 probe sets remained after AYATS quality control, these mapped to only 7,347 unique Entrez IDs. In PREG, the 12,249 remaining probe sets mapped to 8,069 unique genes. Every probe set that mapped to a single gene was retained, including those genes that were measured by more than one probe set (i.e., duplicate Entrez identifiers were retained). Multiple probe sets measuring expression of the same gene could provide additional information about differences in expression among various transcripts.[6]

Following summarization and annotation, non-specific probe sets that mapped to multiple Entrez identifiers were removed in order to avoid inaccurate expression values arising from cross-hybridization. To reduce the impact of confounding by sex, all probe sets located on X and Y chromosomes were removed prior to analysis. Finally, to retain only those genes currently expressed in the samples, the MAS5.0 detection calls algorithm was used to determine whether transcripts were present or absent.[29,30] Genes were considered unexpressed and removed if they received absent calls in more than 20% of the samples.[31]

### 4 Primary Results

After multiple testing correction, a total of 903 associations were identified in the AYATS cohort, and 379 DNAm-GE associations were statistically significant in PREG. Across all categories (i.e., AYATS/PREG *cis*/*trans*), many significant relationships occurred between one transcript and one CpG site, although multiple connections were also common (Figures S1 and S2).

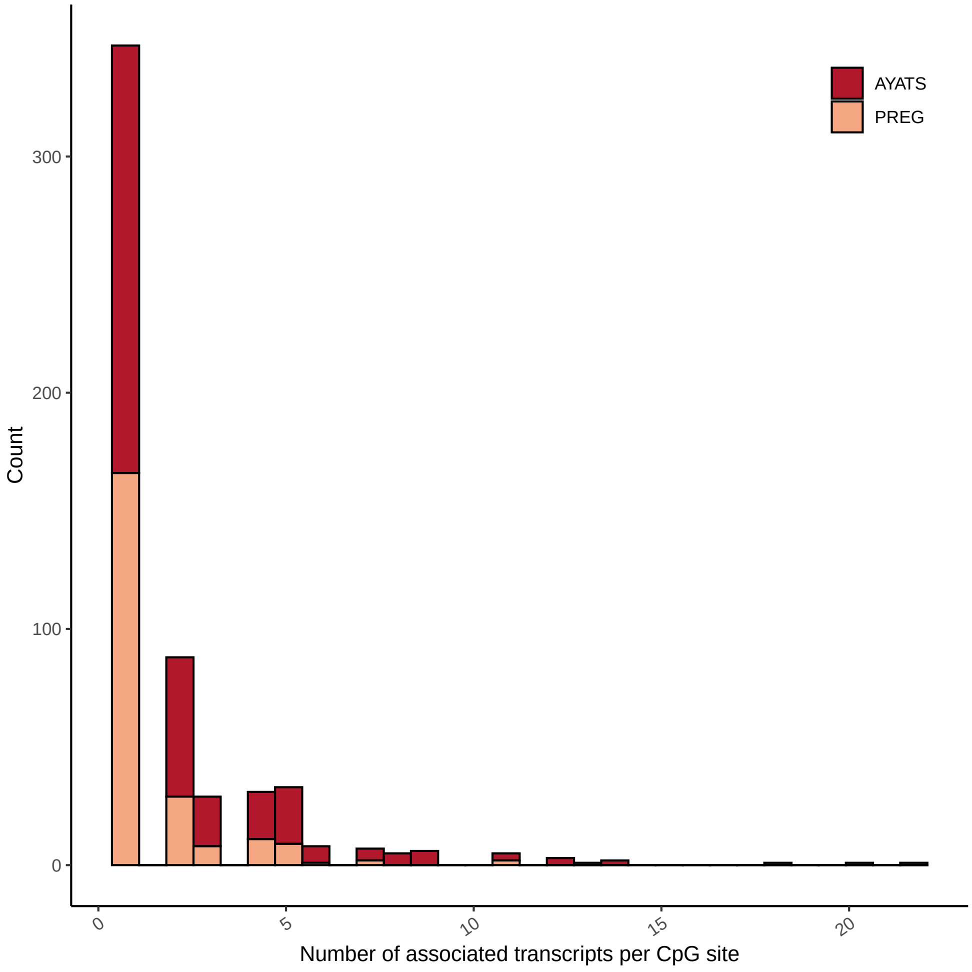

Figure S1: **Distribution of significant CpG associations.** Although PREG had less significant relationships overall, distributions were similar across both the AYATS (red) and PREG (pink) cohorts.

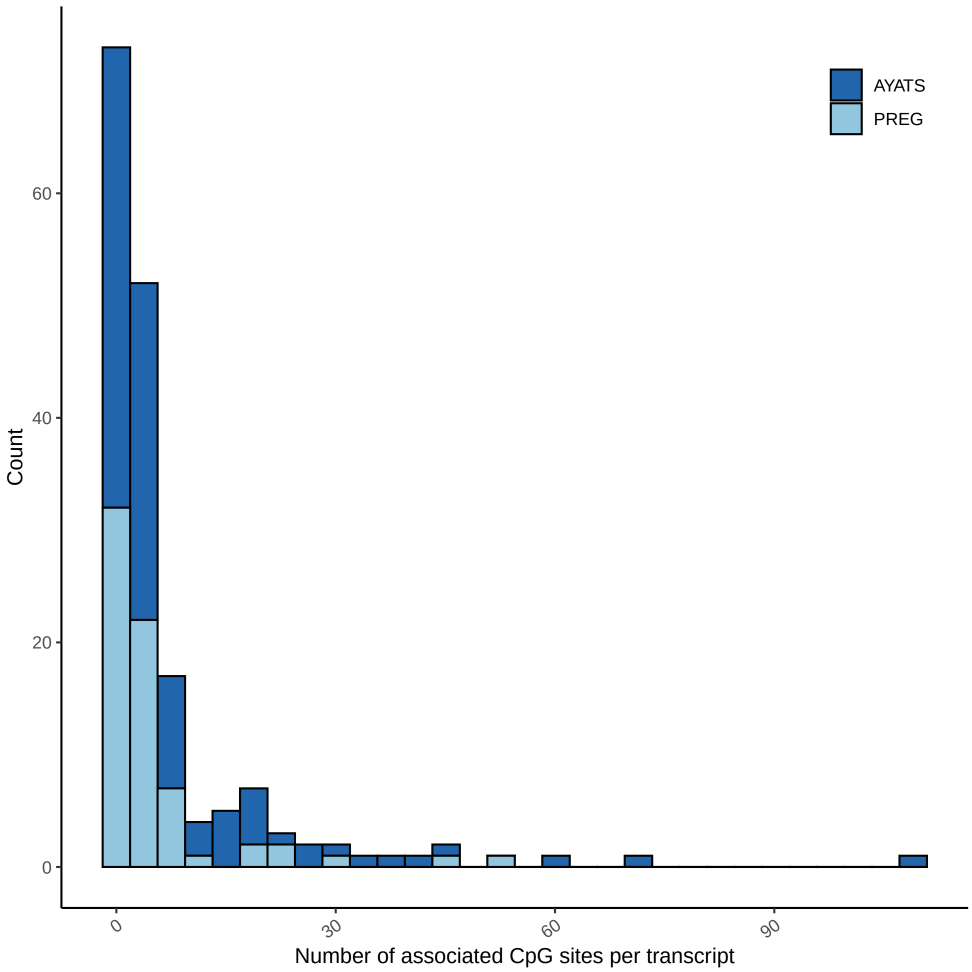

Figure S2: **Distribution of significant transcript associations.** Although PREG had less significant relationships overall, distributions were similar across both the AYATS (dark blue) and PREG (light blue) cohorts.

### 5 Enrichment Analyses

#### 5.1 Local CpG density

To determine the characteristics of CpG sites that associate with transcript expression, all significant CpGs were functionally annotated and tested for enrichment. Both UCSC CpG island classifiers and HIL annotations describe local CpG density. Although CpG island classifiers are more commonly used in the literature, evidence suggests that HIL definitions more accurately describe a distinctive pattern of CpGs.[15]

*Cis* and *trans* results were tested separately to determine characteristics that differ depending on if CpG sites are in a proximal (Figure S3) or distal (Figure S4) relationship with the associated transcript. Regions of low CpG density were enriched in both *cis* and *trans* in AYATS only, while intermediate density regions were enriched in *trans* across both cohorts. High density regions were consistently depleted in AYATS and PREG (Table S3).

Table S3: Results of CpG Density Enrichment Analyses[1]

| Annotation | AYATS | | | | PREG | | | |
|---|---|---|---|---|---|---|---|---|
| | *cis* | | *trans* | | *cis* | | *trans* | |
| | enriched | depleted | enriched | depleted | enriched | depleted | enriched | depleted |
| HIL Classifiers[2] | **LC** | IC shore, <u>**HC**</u> | <u>**IC**</u>, **LC** | ICshore, <u>**HC**</u> | IC | <u>HC</u> | <u>**IC**</u>, IC shore | <u>**HC**</u> |

**Bolded items.** $P$-val $< 0.0025$ (Bonferroni corrected for 20 tests)
<u>Underlined items.</u> Concordance across both the AYATS and PREG study.
[1] $P$-val $< 0.05$
[2] CpG density classifiers; high density (HC), low density (LC), intermediate density (IC), and intermediate density bordering high density regions (IC shore)

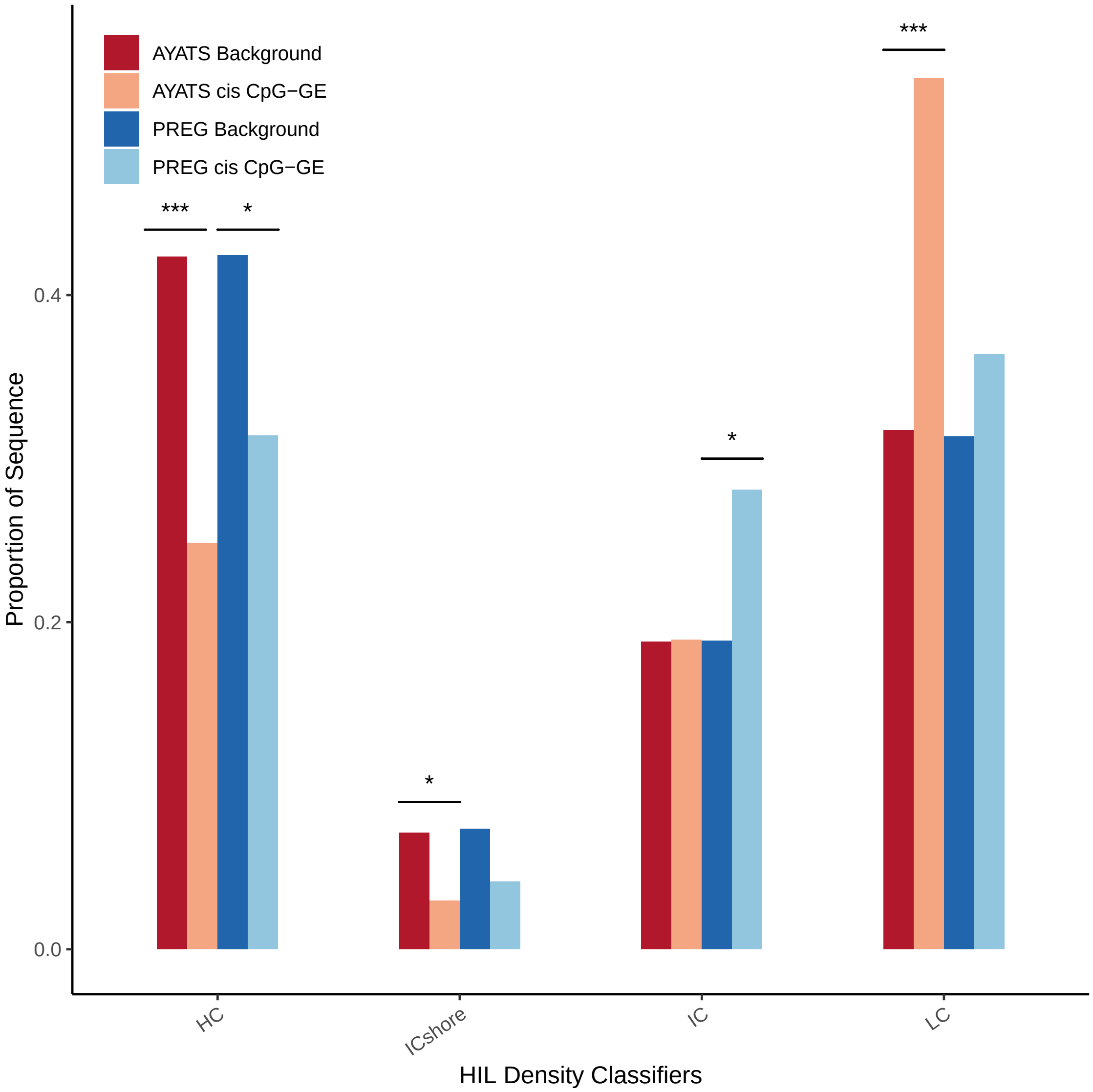

Figure S3: **Enrichment for HIL CpG density classifiers in *cis* CpG-transcript relationships.** Overall, areas of higher CpG density were depleted while areas of intermediate and low CpG density were enriched in GE-associated CpGs (*** = $p < 0.0005$; ** = $p < 0.005$; * = $p < 0.05$).

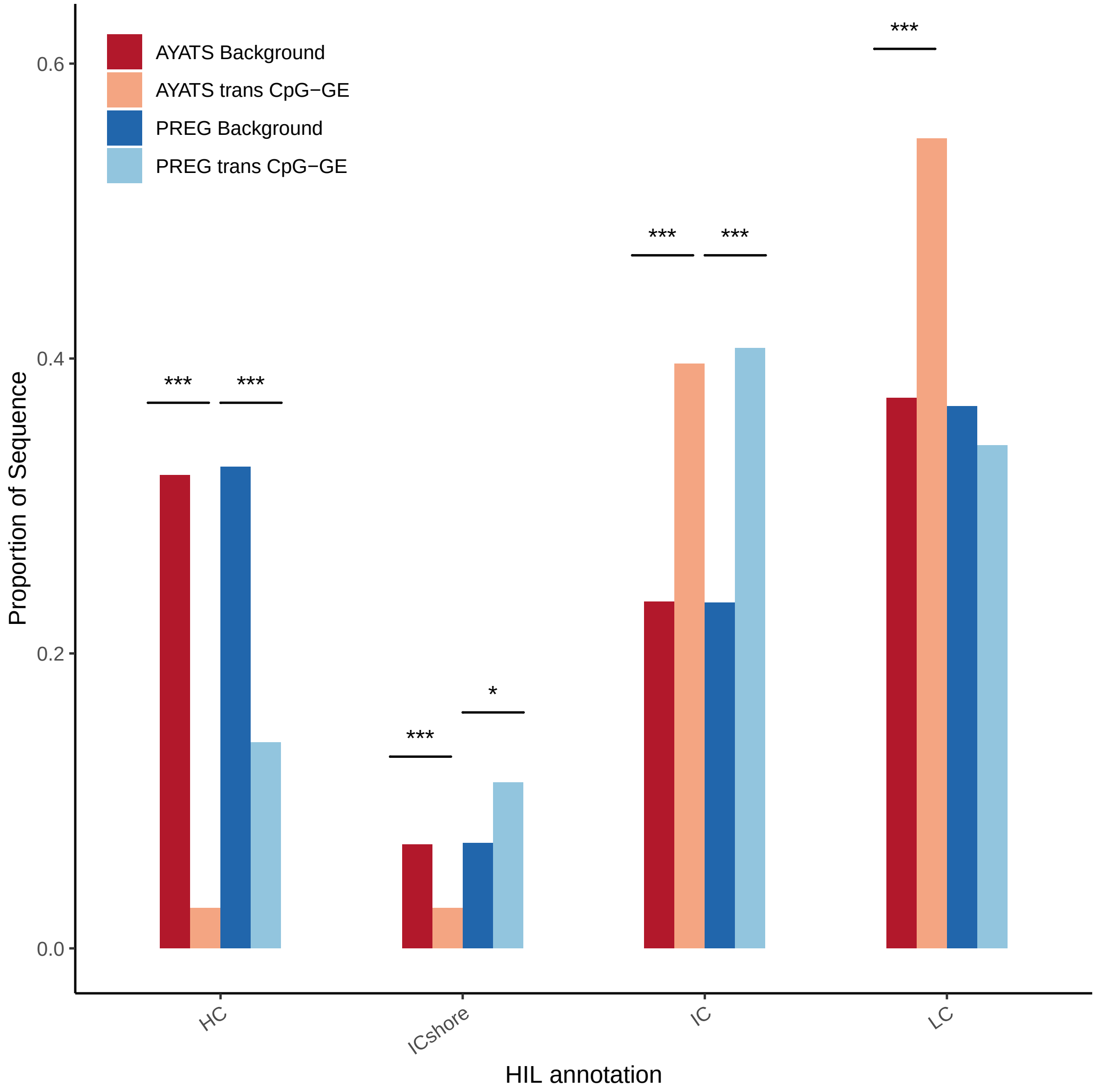

Figure S4: **Enrichment for HIL CpG density classifiers in *trans* CpG-transcript relationships.** High density regions were depleted while intermediate density regions were enriched. The regions bordering these annotations (ICshores) were less consistent across cohorts, showing enrichment in PREG but depletion in AYATS (*** = $p < 0.0005$; ** = $p < 0.005$; * = $p < 0.05$).

##### 5.2 Gene set enrichment analysis

Enrichment for biological process, molecular function, and cellular function gene ontology (GO) terms was examined using the `clusterProfiler` package in R.[32] For a thorough characterization of molecular pathways related to *cis* DNAm-GE relationships, enrichment for KEGG pathways was also examined. GE probe sets had previously been annotated to their respective Entrez identifiers. The distribution of all tested *cis* transcripts was compared to those significantly associated with DNAm at one or more *cis* CpGs. Duplicates were conserved, so that in cases where a transcript was tested against several *cis* CpGs the transcript was repeated accordingly. This detail is important for controlling for the background bias of the microarray. Terms with a false discovery rate (FDR) $<$ 0.05 were considered significant.[33]

Common themes were uncovered in both cohorts (Tables S4-S9). These include functions related to the activation and regulation of immune response, and those related to adaptive immunity. Terms related to cellular detoxification and control of the cellular response to outside elements were also enriched within significant results. Further, vesicle and lysosome formation and transport were represented in molecular function analyses. A total of 33 terms overlapped between the two cohorts, with 78.6% of PREG terms also found in AYATS.

Enrichment for KEGG pathways were examined using the same methods described above. Again, significant pathways tended to overlap between the two cohorts. All significant pathways identified in AYATS (n = 31) were also found in PREG (n = 39). Pathways related to infectious and autoimmune diseases were heavily represented (Tables S10-S11).

Although many overlapping processes and pathways were enriched across cohorts, it remains important to note that a relatively few number of genes were implicated in the majority of replicated terms.

Table S4: AYATS Gene Ontology Biological Processes Enrichment Results[1]

| Term Identifier[2] | Term Name | q-value | Genes |
|---|---|---|---|
| **GO:1901685** | glutathione derivative metabolic process | 0.0080 | GSTM3/MGST3/GSTM1 |
| **GO:1901687** | glutathione derivative biosynthetic process | 0.0080 | GSTM3/MGST3/GSTM1 |
| GO:0002768 | immune response-regulating cell surface receptor signaling pathway | 0.0225 | HLA-DPA1/HLA-DQB1/BTNL8/SOS1/HLA-DQA1/ C3AR1/BTN3A2 |
| GO:0002250 | adaptive immune response | 0.0437 | HLA-DPA1/HLA-DQB1/BTNL8/HLA-DQA1/ IL18RAP/BTN3A2 |
| GO:0050852 | T cell receptor signaling pathway | 0.0437 | HLA-DPA1/HLA-DQB1/BTNL8/HLA-DQA1/ BTN3A2 |

[1] $q < 0.05$
[2] Bolded identifiers represent those terms significant in both the AYATS and PREG cohorts.

Table S5: AYATS Gene Ontology Molecular Function Enrichment Results[1]

| Term Identifier[2] | Term Name | q-value | Genes |
|---|---|---|---|
| **GO:0004364** | glutathione transferase activity | 0.0020 | GSTM3/MGST3/GSTM1 |
| **GO:0042605** | peptide antigen binding | 0.0022 | HLA-DPA1/HLA-DQB1/HLA-DQA1 |
| **GO:0042277** | peptide binding | 0.0064 | HLA-DPA1/HLA-DQB1/HLA-DQA1/GSTM3/ GSTM1 |
| **GO:0003823** | antigen binding | 0.0068 | HLA-DPA1/HLA-DQB1/HLA-DQA1 |
| **GO:0016765** | transferase activity, transferring alkyl or aryl (other than methyl) groups | 0.0068 | GSTM3/MGST3/GSTM1 |
| **GO:0033218** | amide binding | 0.0084 | HLA-DPA1/HLA-DQB1/HLA-DQA1/GSTM3/ GSTM1 |
| GO:0004888 | transmembrane signaling receptor activity | 0.0460 | HLA-DPA1/HLA-DQB1/HLA-DQA1/IL18RAP/ C3AR1 |

[1] $q < 0.05$
[2] Bolded identifiers represent those terms significant in both the AYATS and PREG cohorts.

Table S6: AYATS Gene Ontology Cellular Components Enrichment Results[1]

| Term Identifier[2] | Term Name | q-value | Genes |
|---|---|---|---|
| **GO:0071556** | integral component of lumenal side of endoplasmic reticulum membrane | 0.0015 | HLA-DPA1/HLA-DQB1/HLA-DQA1 |
| **GO:0098553** | lumenal side of endoplasmic reticulum membrane | 0.0015 | HLA-DPA1/HLA-DQB1/HLA-DQA1 |
| **GO:0042611** | MHC protein complex | 0.0015 | HLA-DPA1/HLA-DQB1/HLA-DQA1 |
| **GO:0030669** | clathrin-coated endocytic vesicle membrane | 0.0019 | HLA-DPA1/HLA-DQB1/HLA-DQA1 |
| **GO:0000323** | lytic vacuole | 0.0030 | HLA-DPA1/HLA-DQB1/HLA-DQA1/CTSW/C3AR1/ AGA/LIPA/RNASET2 |
| **GO:0005764** | lysosome | 0.0030 | HLA-DPA1/HLA-DQB1/HLA-DQA1/CTSW/C3AR1/ AGA/LIPA/RNASET2 |
| **GO:0045334** | clathrin-coated endocytic vesicle | 0.0030 | HLA-DPA1/HLA-DQB1/HLA-DQA1 |
| **GO:0044437** | vacuolar part | 0.0047 | HLA-DPA1/HLA-DQB1/HLA-DQA1/C3AR1/AGA/ LIPA/RNASET2 |
| **GO:0005773** | vacuole | 0.0050 | HLA-DPA1/HLA-DQB1/HLA-DQA1/CTSW/C3AR1/ AGA/LIPA/RNASET2 |
| **GO:0012507** | ER to Golgi transport vesicle membrane | 0.0075 | HLA-DPA1/HLA-DQB1/HLA-DQA1 |
| **GO:0032588** | trans-Golgi network membrane | 0.0083 | HLA-DPA1/HLA-DQB1/HLA-DQA1 |
| **GO:0030665** | clathrin-coated vesicle membrane | 0.0128 | HLA-DPA1/HLA-DQB1/HLA-DQA1 |
| **GO:0030134** | COPII-coated ER to Golgi transport vesicle | 0.0167 | HLA-DPA1/HLA-DQB1/HLA-DQA1 |
| **GO:0030660** | Golgi-associated vesicle membrane | 0.0228 | HLA-DPA1/HLA-DQB1/HLA-DQA1 |
| GO:0098552 | side of membrane | 0.0246 | HLA-DPA1/HLA-DQB1/BTNL8/HLA-DQA1/BTN3A2 |
| **GO:0030666** | endocytic vesicle membrane | 0.0361 | HLA-DPA1/HLA-DQB1/HLA-DQA1 |
| **GO:0030176** | integral component of endoplasmic reticulum membrane | 0.0361 | HLA-DPA1/HLA-DQB1/HLA-DQA1 |
| **GO:0031227** | intrinsic component of endoplasmic reticulum membrane | 0.0361 | HLA-DPA1/HLA-DQB1/HLA-DQA1 |
| **GO:0030136** | clathrin-coated vesicle | 0.0361 | HLA-DPA1/HLA-DQB1/HLA-DQA1 |
| GO:0005775 | vacuolar lumen | 0.0361 | AGA/LIPA/RNASET2 |
| **GO:0030658** | transport vesicle membrane | 0.0370 | HLA-DPA1/HLA-DQB1/HLA-DQA1 |
| **GO:0030662** | coated vesicle membrane | 0.0370 | HLA-DPA1/HLA-DQB1/HLA-DQA1 |
| **GO:0045171** | intercellular bridge | 0.0370 | GSTM3/GSTM1 |
| GO:0005766 | primary lysosome | 0.0370 | C3AR1/AGA/RNASET2 |
| GO:0042582 | azurophil granule | 0.0370 | C3AR1/AGA/RNASET2 |
| **GO:0005798** | Golgi-associated vesicle | 0.0384 | HLA-DPA1/HLA-DQB1/HLA-DQA1 |
| GO:0034774 | secretory granule lumen | 0.0411 | CTSW/AGA/CHI3L1/RNASET2 |
| **GO:0005765** | lysosomal membrane | 0.0411 | HLA-DPA1/HLA-DQB1/HLA-DQA1/C3AR1 |
| **GO:0098852** | lytic vacuole membrane | 0.0411 | HLA-DPA1/HLA-DQB1/HLA-DQA1/C3AR1 |
| GO:0098797 | plasma membrane protein complex | 0.0411 | HLA-DPA1/HLA-DQB1/HLA-DQA1/IL18RAP |
| GO:0060205 | cytoplasmic vesicle lumen | 0.0416 | CTSW/AGA/CHI3L1/RNASET2 |
| GO:0031983 | vesicle lumen | 0.0416 | CTSW/AGA/CHI3L1/RNASET2 |
| **GO:0005802** | trans-Golgi network | 0.0450 | HLA-DPA1/HLA-DQB1/HLA-DQA1 |
| GO:0043202 | lysosomal lumen | 0.0459 | LIPA/RNASET2 |
| GO:0030141 | secretory granule | 0.0459 | CTSW/RNPEP/C3AR1/AGA/CHI3L1/RNASET2 |

[1] $q < 0.05$
[2] Bolded identifiers represent those terms significant in both the AYATS and PREG cohorts.

Table S7: PREG Gene Ontology Biological Processes Enrichment Results[1]

| Term Identifier[2] | Term Name | q-value | Genes |
|---|---|---|---|
| **GO:1901685** | glutathione derivative metabolic process | 0.0065 | GSTT1/GSTM3/GSTM1 |
| **GO:1901687** | glutathione derivative biosynthetic process | 0.0065 | GSTT1/GSTM3/GSTM1 |
| GO:0006805 | xenobiotic metabolic process | 0.0065 | CES1/GSTM3/SULT1A2/GSTM1 |
| GO:0071466 | cellular response to xenobiotic stimulus | 0.0275 | CES1/GSTM3/SULT1A2/GSTM1 |
| GO:0006749 | glutathione metabolic process | 0.0275 | GSTT1/GSTM3/GSTM1 |

[1] $q < 0.05$

[2] Bolded identifiers represent those terms significant in both the AYATS and PREG cohorts.

Table S8: PREG Gene Ontology Molecular Function Enrichment Results[1]

| Term Identifier[2] | Term Name | q-value | Genes |
|---|---|---|---|
| **GO:0004364** | glutathione transferase activity | 0.0026 | GSTT1/GSTM3/GSTM1 |
| **GO:0042605** | peptide antigen binding | 0.0026 | HLA-DQA1/HLA-DQB1/HLA-DPA1 |
| **GO:0042277** | peptide binding | 0.0092 | HLA-DQA1/HLA-DQB1/HLA-DPA1/GSTM3/GSTM1 |
| **GO:0003823** | antigen binding | 0.0092 | HLA-DQA1/HLA-DQB1/HLA-DPA1 |
| **GO:0016765** | transferase activity, transferring alkyl or aryl (other than methyl) groups | 0.0092 | GSTT1/GSTM3/GSTM1 |
| GO:0016298 | lipase activity | 0.0133 | CES1/LIPA/PGAP6 |
| **GO:0033218** | amide binding | 0.0138 | HLA-DQA1/HLA-DQB1/HLA-DPA1/GSTM3/GSTM1 |
| GO:0052689 | carboxylic ester hydrolase activity | 0.0157 | CES1/LIPA/PGAP6 |

[1] $q < 0.05$
[2] Bolded identifiers represent those terms significant in both the AYATS and PREG cohorts.

Table S9: PREG Gene Ontology Cellular Components Enrichment Results[1]

| Term Identifier[2] | Term Name | q-value | Genes |
|---|---|---|---|
| GO:0042613 | MHC class II protein complex | 0.0004 | HLA-DQA1/HLA-DQB1/HLA-DPA1 |
| **GO:0071556** | integral component of lumenal side of endoplasmic reticulum membrane | 0.0006 | HLA-DQA1/HLA-DQB1/HLA-DPA1 |
| **GO:0098553** | lumenal side of endoplasmic reticulum membrane | 0.0006 | HLA-DQA1/HLA-DQB1/HLA-DPA1 |
| **GO:0042611** | MHC protein complex | 0.0006 | HLA-DQA1/HLA-DQB1/HLA-DPA1 |
| **GO:0030669** | clathrin-coated endocytic vesicle membrane | 0.0012 | HLA-DQA1/HLA-DQB1/HLA-DPA1 |
| **GO:0045334** | clathrin-coated endocytic vesicle | 0.0030 | HLA-DQA1/HLA-DQB1/HLA-DPA1 |
| **GO:0012507** | ER to Golgi transport vesicle membrane | 0.0060 | HLA-DQA1/HLA-DQB1/HLA-DPA1 |
| GO:0005865 | striated muscle thin filament | 0.0060 | TNNT1/MYOM2 |
| GO:0036379 | myofilament | 0.0060 | TNNT1/MYOM2 |
| **GO:0032588** | trans-Golgi network membrane | 0.0060 | HLA-DQA1/HLA-DQB1/HLA-DPA1 |
| **GO:0030665** | clathrin-coated vesicle membrane | 0.0096 | HLA-DQA1/HLA-DQB1/HLA-DPA1 |
| **GO:0005773** | vacuole | 0.0096 | HLA-DQA1/CTSW/HLA-DQB1/HLA-DPA1/GLB1L/ LIPA/PGAP6 |
| **GO:0030134** | COPII-coated ER to Golgi transport vesicle | 0.0101 | HLA-DQA1/HLA-DQB1/HLA-DPA1 |
| **GO:0030660** | Golgi-associated vesicle membrane | 0.0140 | HLA-DQA1/HLA-DQB1/HLA-DPA1 |
| **GO:0000323** | lytic vacuole | 0.0194 | HLA-DQA1/CTSW/HLA-DQB1/HLA-DPA1/LIPA/ PGAP6 |
| **GO:0005764** | lysosome | 0.0194 | HLA-DQA1/CTSW/HLA-DQB1/HLA-DPA1/LIPA/ PGAP6 |
| **GO:0030176** | integral component of endoplasmic reticulum membrane | 0.0209 | HLA-DQA1/HLA-DQB1/HLA-DPA1 |
| **GO:0030666** | endocytic vesicle membrane | 0.0209 | HLA-DQA1/HLA-DQB1/HLA-DPA1 |
| **GO:0031227** | intrinsic component of endoplasmic reticulum membrane | 0.0209 | HLA-DQA1/HLA-DQB1/HLA-DPA1 |
| **GO:0030136** | clathrin-coated vesicle | 0.0227 | HLA-DQA1/HLA-DQB1/HLA-DPA1 |
| **GO:0030658** | transport vesicle membrane | 0.0249 | HLA-DQA1/HLA-DQB1/HLA-DPA1 |
| **GO:0030662** | coated vesicle membrane | 0.0249 | HLA-DQA1/HLA-DQB1/HLA-DPA1 |
| **GO:0005798** | Golgi-associated vesicle | 0.0249 | HLA-DQA1/HLA-DQB1/HLA-DPA1 |
| **GO:0044437** | vacuolar part | 0.0249 | HLA-DQA1/HLA-DQB1/HLA-DPA1/LIPA/PGAP6 |
| **GO:0005765** | lysosomal membrane | 0.0249 | HLA-DQA1/HLA-DQB1/HLA-DPA1/PGAP6 |
| **GO:0098852** | lytic vacuole membrane | 0.0249 | HLA-DQA1/HLA-DQB1/HLA-DPA1/PGAP6 |
| **GO:0045171** | intercellular bridge | 0.0249 | GSTM3/GSTM1 |
| **GO:0005802** | trans-Golgi network | 0.0337 | HLA-DQA1/HLA-DQB1/HLA-DPA1 |
| GO:0005774 | vacuolar membrane | 0.0352 | HLA-DQA1/HLA-DQB1/HLA-DPA1/PGAP6 |

[1] $q < 0.05$

[2] Bolded identifiers represent those terms significant in both the AYATS and PREG cohorts.

Table S10: AYATS Enrichment of KEGG Pathways Results[1]

| Term Identifier[2] | Term Name | q-value | Entrez |
|---|---|---|---|
| **hsa05150** | Staphylococcus aureus infection | 0.0006 | 3113/3119/3117/719 |
| **hsa05310** | Asthma | 0.0006 | 3113/3119/3117 |
| **hsa05321** | Inflammatory bowel disease (IBD) | 0.0007 | 3113/3119/3117/8807 |
| **hsa00982** | Drug metabolism - cytochrome P450 | 0.0017 | 2947/4259/2944 |
| **hsa05320** | Autoimmune thyroid disease | 0.0017 | 3113/3119/3117 |
| **hsa00980** | Metabolism of xenobiotics by cytochrome P450 | 0.0017 | 2947/4259/2944 |
| **hsa05330** | Allograft rejection | 0.0017 | 3113/3119/3117 |
| **hsa04672** | Intestinal immune network for IgA production | 0.0017 | 3113/3119/3117 |
| **hsa05332** | Graft-versus-host disease | 0.0017 | 3113/3119/3117 |
| **hsa04940** | Type I diabetes mellitus | 0.0017 | 3113/3119/3117 |
| **hsa05204** | Chemical carcinogenesis | 0.0017 | 2947/4259/2944 |
| **hsa00480** | Glutathione metabolism | 0.0024 | 2947/4259/2944 |
| **hsa00983** | Drug metabolism - other enzymes | 0.0035 | 2947/4259/2944 |
| **hsa05416** | Viral myocarditis | 0.0041 | 3113/3119/3117 |
| **hsa05225** | Hepatocellular carcinoma | 0.0048 | 6654/2947/4259/2944 |
| **hsa05322** | Systemic lupus erythematosus | 0.0048 | 3113/3119/3117 |
| **hsa04612** | Antigen processing and presentation | 0.0058 | 3113/3119/3117 |
| **hsa05140** | Leishmaniasis | 0.0062 | 3113/3119/3117 |
| **hsa01524** | Platinum drug resistance | 0.0078 | 2947/4259/2944 |
| **hsa05323** | Rheumatoid arthritis | 0.0082 | 3113/3119/3117 |
| **hsa04640** | Hematopoietic cell lineage | 0.0100 | 3113/3119/3117 |
| **hsa04658** | Th1 and Th2 cell differentiation | 0.0100 | 3113/3119/3117 |
| **hsa04514** | Cell adhesion molecules (CAMs) | 0.0109 | 3113/3119/3117 |
| **hsa05145** | Toxoplasmosis | 0.0129 | 3113/3119/3117 |
| **hsa05166** | Human T-cell leukemia virus 1 infection | 0.0143 | 3113/3119/3117/8379 |
| **hsa04659** | Th17 cell differentiation | 0.0156 | 3113/3119/3117 |
| **hsa05418** | Fluid shear stress and atherosclerosis | 0.0224 | 2947/4259/2944 |
| **hsa04142** | Lysosome | 0.0224 | 1521/175/3988 |
| **hsa04145** | Phagosome | 0.0304 | 3113/3119/3117 |
| **hsa05164** | Influenza A | 0.0360 | 3113/3119/3117 |
| **hsa05152** | Tuberculosis | 0.0401 | 3113/3119/3117 |

[1] $q < 0.05$

[2] Bolded identifiers represent those terms significant in both the AYATS and PREG cohorts.

Table S11: PREG Enrichment of KEGG Pathways Results[1]

| Term Identifier[2] | Term Name | q-value | Entrez |
|---|---|---|---|
| **hsa05204** | Chemical carcinogenesis | 0.0001 | 2952/2947/6799/2944 |
| **hsa00983** | Drug metabolism - other enzymes | 0.0001 | 2952/1066/2947/2944 |
| **hsa05310** | Asthma | 0.0001 | 3117/3119/3113 |
| **hsa00982** | Drug metabolism - cytochrome P450 | 0.0003 | 2952/2947/2944 |
| **hsa05320** | Autoimmune thyroid disease | 0.0003 | 3117/3119/3113 |
| **hsa00980** | Metabolism of xenobiotics by cytochrome P450 | 0.0003 | 2952/2947/2944 |
| **hsa05330** | Allograft rejection | 0.0003 | 3117/3119/3113 |
| **hsa04672** | Intestinal immune network for IgA production | 0.0003 | 3117/3119/3113 |
| **hsa05332** | Graft-versus-host disease | 0.0003 | 3117/3119/3113 |
| **hsa04940** | Type I diabetes mellitus | 0.0003 | 3117/3119/3113 |
| **hsa00480** | Glutathione metabolism | 0.0003 | 2952/2947/2944 |
| **hsa05150** | Staphylococcus aureus infection | 0.0005 | 3117/3119/3113 |
| **hsa05416** | Viral myocarditis | 0.0006 | 3117/3119/3113 |
| **hsa05321** | Inflammatory bowel disease (IBD) | 0.0006 | 3117/3119/3113 |
| **hsa04612** | Antigen processing and presentation | 0.0009 | 3117/3119/3113 |
| **hsa05140** | Leishmaniasis | 0.0010 | 3117/3119/3113 |
| **hsa01524** | Platinum drug resistance | 0.0011 | 2952/2947/2944 |
| **hsa05322** | Systemic lupus erythematosus | 0.0011 | 3117/3119/3113 |
| **hsa05323** | Rheumatoid arthritis | 0.0011 | 3117/3119/3113 |
| **hsa04658** | Th1 and Th2 cell differentiation | 0.0014 | 3117/3119/3113 |
| **hsa04640** | Hematopoietic cell lineage | 0.0015 | 3117/3119/3113 |
| **hsa04514** | Cell adhesion molecules (CAMs) | 0.0015 | 3117/3119/3113 |
| **hsa05145** | Toxoplasmosis | 0.0020 | 3117/3119/3113 |
| **hsa04659** | Th17 cell differentiation | 0.0021 | 3117/3119/3113 |
| **hsa05418** | Fluid shear stress and atherosclerosis | 0.0032 | 2952/2947/2944 |
| **hsa05225** | Hepatocellular carcinoma | 0.0042 | 2952/2947/2944 |
| **hsa04145** | Phagosome | 0.0045 | 3117/3119/3113 |
| **hsa05164** | Influenza A | 0.0056 | 3117/3119/3113 |
| **hsa05152** | Tuberculosis | 0.0061 | 3117/3119/3113 |
| hsa05169 | Epstein-Barr virus infection | 0.0110 | 3117/3119/3113 |
| **hsa05166** | Human T-cell leukemia virus 1 infection | 0.0114 | 3117/3119/3113 |
| hsa00100 | Steroid biosynthesis | 0.0204 | 3988 |
| **hsa04142** | Lysosome | 0.0208 | 1521/3988 |
| hsa05168 | Herpes simplex virus 1 infection | 0.0242 | 3117/3119/3113 |
| hsa04966 | Collecting duct acid secretion | 0.0255 | 10723 |
| hsa04979 | Cholesterol metabolism | 0.0301 | 3988 |
| hsa04978 | Mineral absorption | 0.0331 | 55630 |
| hsa04974 | Protein digestion and absorption | 0.0383 | 80781 |
| hsa05200 | Pathways in cancer | 0.0447 | 2952/2947/2944 |

[1] $q < 0.05$

[2] Bolded identifiers represent those terms significant in both the AYATS and PREG cohorts.

[6] *Technical Note, Design and Performance of the GeneChip® Human Genome U133 Plus 2.0 and Human Genome U133A 2.0 Array*, 2003.

[7] R Core Team, "R: A language and environment for statistical computing. R Foundation for Statistical Computing, Vienna, Austria.," 2016.

[21] A. Wolen, *mimager: The Microarray Imager*, 2019. R package version 1.10.0.

[22] R. T. Barfield, L. M. Almli, V. Kilaru, A. K. Smith, K. B. Mercer, R. Duncan, T. Klengel, D. Mehta, E. B. Binder, M. P. Epstein, K. J. Ressler, and K. N. Conneely, “Accounting for population stratification in DNA methylation studies,” *Genet. Epidemiol.*, vol. 38, pp. 231–241, Apr. 2014.

[23] M. N. McCall, P. N. Murakami, M. Lukk, W. Huber, and R. A. Irizarry, “Assessing Affymetrix GeneChip microarray quality,” *BMC Bioinformatics*, vol. 12, p. 137, May 2011.

[24] R. A. Irizarry, B. Hobbs, F. Collin, Y. D. Beazer-Barclay, K. J. Antonellis, U. Scherf, and T. P. Speed, “Exploration, normalization, and summaries of high density oligonucleotide array probe level data,” *Biostatistics*, vol. 4, pp. 249–264, Apr. 2003.

[25] J. T. Leek, W. E. Johnson, H. S. Parker, E. J. Fertig, A. E. Jaffe, J. D. Storey, Y. Zhang, and L. C. Torres, *sva: Surrogate Variable Analysis*, 2019. R package version 3.34.0.

[26] M. Carlson and B. P. Maintainer, *TxDb.Hsapiens.UCSC.hg19.knownGene: Annotation package for TxDb object(s)*, 2015. R package version 3.2.2.
